# TSPAN10 Loss-of-Function in hESC-Derived RPE Reveals Critical Roles in Pigmentation, Polarity, and Antioxidant Defence

**DOI:** 10.64898/2026.11.19.760930

**Authors:** Rebecca Kaye, Liliya Nazlamova, Eloise Keeling, Magdalena Martinez, Jörn Lakowski, Andrew J Lotery

## Abstract

Expression quantitative trait loci studies have highlighted *TSPAN10* as a gene that confers an increased risk of developing age-related macular degeneration (AMD). The mechanism for this and the role of TSPAN10 in retinal cells such as the retinal pigment epithelium (RPE) and its contribution to AMD pathology are unknown. We report the generation of human embryonic stem cell derived RPE with TSPAN10 knockout using CRISPR/Cas9 genome editing. This provided an in vitro model to investigate the functions of TSPAN10. TSPAN10 knockout cells were differentiated into mature RPE. Significant differences were observed in cell pigmentation, polarisation and cell body height, and the response to oxidative stress in comparison to wild type. These results demonstrate for the first time a role for TSPAN10 in the RPE, aligning with published data to support the importance of TSPAN10 for pigmentation, and adding to our understanding of the heterogeneous disease AMD.

## Introduction

Age related macular degeneration (AMD) is the leading cause of vision loss in the Western World ^1^, accounting for 5.6% of global blindness^1^. Irreversible visual impairment arises due to the gradual loss of photoreceptors, and their supportive cells: the retinal pigment epithelium (RPE). Currently therapeutics are only available to treat approximately 50% of patients, primarily those suffering from the neovascular (wet) form of the disease, leaving a significant unmet clinical need for those progressing to geographic atrophy, late-stage dry AMD. This is due to the heterogeneity of disease and an incomplete understanding of the pathophysiology. Genetic risk is a key element of AMD susceptibility, and multiple risk loci have been identified through Genome Wide Association Studies (GWAS) including a novel polymorphism at *NPLOC4/Tetraspanin 10* (*TSPAN10*) ^2,3^. More recently other genomic methods complementing GWAS have been employed in AMD including expression quantitative trait loci studies (eQTL) ^3^. eQTL integrate genomic and transcriptomic data sets to confirm whether causal genes found in GWAS studies are driving disease association. In particular a recent eQTL study highlighted *TSPAN10* at the *NPLOC4/TSPAN10* locus as a significant target gene associated with AMD ^3^.

The *TSPAN10* single nucleotide polymorphism (SNP) associated with an increased risk of AMD is rs6565597, OR 1.13 ^2^.^3^. eQTL data from donor eyes has identified reduced copy numbers of *TSPAN10* in those heterozygous/homozygous for rs6565597 ^3^. Recently the rs6565597 variant has again been associated with late AMD, this time in an older age group, specifically those over 90 years, OR 1.46 ^4^. Interestingly we have previously shown rs6565597 to be significantly associated with outer retinal thinning at the macula in clinically normal individuals in the UK Biobank using automated layer segmentation ^5^. Changes were seen specifically between the photoreceptor inner-segment outer-segment to RPE (ISOS-RPE) and RPE to Bruch’s membrane (RPE-BM)^6^. Despite this association, the function of TSPAN10 in the retina is unknown.

Tetraspanins are a large family of integral membrane proteins typically located at the plasma membrane. TSPAN10 belongs to the tetraspanin family, specifically, the C8 tetraspanins, also known as TspanC8. All members of the TspanC8 subgroup regulate the exit of the transmembrane matrix metalloprotease ADAM10 from the endoplasmic reticulum (ER), and the majority also regulate its’ maturation ^7^ ^8^. TSPAN10 is also known as Oculospanin given that it was originally discovered in the eye ^9^. It was initially detected in the RPE, iris and ciliary body ^9^. Single-cell RNA sequencing data from the retina and choroid has further isolated TSPAN10 to the RPE and melanocytes within the choroid ^10^.

Within the eye, polymorphisms at *TSPAN10* are associated with myopia ^11^, astigmatism ^11^, strabismus ^12^ and as mentioned an increased risk of AMD ^2,4^. *TSPAN10* SNPs are also significantly associated with the thickness of retinal layers, with reduced thickness at the foveola and a reduction in the thickness of the retinal nerve fibre layer and ganglion cell layer at the macula ^13^. Recently TSPAN10 has been significantly associated with foveal morphology ^14^, suggesting a role in shaping the fovea. The same publication also showed a link between a larger foveal pit volume and an increased future risk of developing AMD.

TSPAN10 is thought to play a role in pigmentation, given its association with lighter hair colour ^15,16^ and role in eye colour ^17,18^. A genetic locus at TSPAN10 has also been associated with the ‘Retinal Pigment Score’ ^19^, a quantitative measure of pigmentation in a colour fundus photograph, with polymorphisms at TSPAN10 associated with a less pigmented fundus.

The function of TSPAN10 in both the eye and specifically the RPE, however, is unknown. Given the relationship TSPAN10 has with AMD risk and macula thinning, we aimed to elucidate its role in the RPE by knocking out the *TSPAN10* gene using the CRISPR/Cas9 system in human embryonic stem cells (hESCs). TSPAN10 deficient hESCs were generated and induced to differentiate into hESC-RPE cells, where the function of TSPAN10 was analysed.

## Results

### Development of TSPAN10^-/-^ hESC line and RPE differentiation

To generate a TSPAN10 knockout (^-/-^) hESC line we aimed to produce a frame-shift mutation through introduction of an indel at the ATG in exon 2 of *TSPAN10* in the MasterShef7 embryonic stem cell line, using CRISPR/Cas9 mediated gene-editing. ^20^ An in-depth description of this approach is provided in the Material and Methods section and is briefly summarised here and in Figure 1 A and Supplementary Figure S1. The plasmid px459 was used to generate biallelic indel mutations in *TSPAN10* with a CRISPR strategy which utilised the non-homologous end joining (NHEJ) pathway. Two clones were isolated containing indels, A2 a 7-base pair (b.p) deletion and A12 a 1 b.p insertion (Figure 1 A), both were predicted to cause a frame shift mutation. To ensure that no off-target mutations were introduced, the top predicted off-sites (crispr.mit.edu) for the sgRNAs were amplified and sequenced from the selected clones and no off-target mutations were detected (Supplemental Table 1, Supplemental Figure S2). Lastly, we performed array comparative genome hybridization (array-CGH) and found that gene editing did not result in the introduction of any copy number variants (Supplemental Figures S3 & S4). All subsequent experiments were conducted using these two homozygous clones (clones A2 and A12).

**Figure 1.**
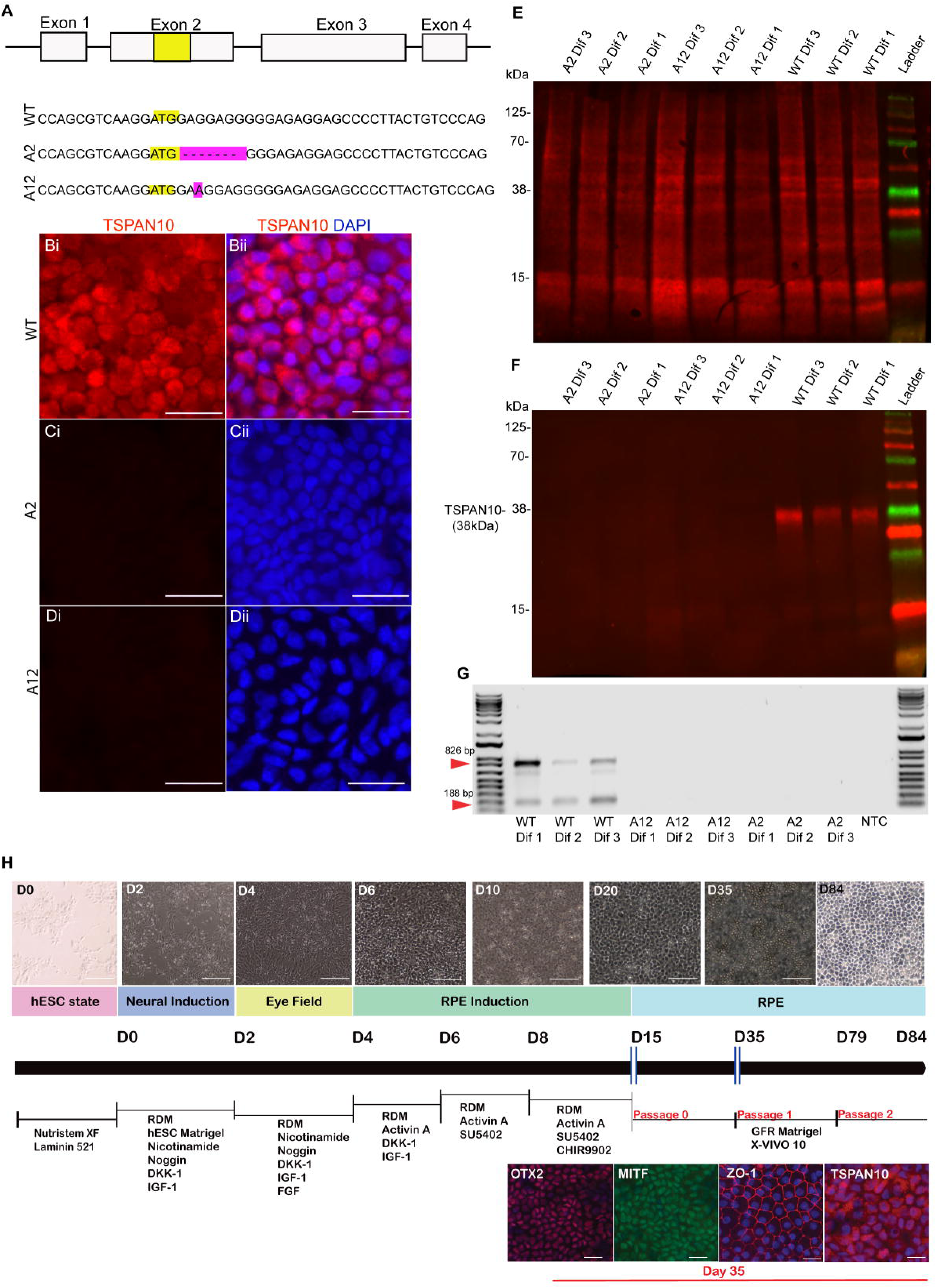
Creation of TSPAN10^-/-^ hESCs using CRISPR/Cas9 and confirmation of TSPAN10^-/-^. **(A)Schematic demonstrating genomic DNA insertions/deletions present in hESC clones.** WT sequence with ATG region highlighted. A2 clone sequence with 7 bp deletion emphasised in purple. A12 clone sequence with insertion of A highlighted in purple. **(B), (C) & (D) Immunocytochemistry of Day 35 hESC-RPE for TSPAN10** Bi, Ci, Di Representative staining of TSPAN10 (red). WT cells display strong expression of TSPAN10. Bii, Cii, Dii Merged images showing dual DAPI and TSPAN10 staining. TSPAN10^-/-^ cell clones A2 & A12 show only DAPI staining with no expression of TSPAN10. Scale bars represent 25µm. Immunocytochemistry performed for three differentiations from all three clones (n=9). (**E) & (F) Western Blot for TSPAN10 protein in *TSPAN10* mutant and WT hESC-RPE cells** E, A total protein stain (Revert 700) was performed as a control to ensure equal loading of lysates. **F,** TSPAN10 protein blot using the Proteintech polyclonal Ab indicates that both clones and all differentiations are true knockouts. N= 3 technical repeats performed **(G) RT-PCR for TSPAN10** in hESC-RPE. Expected product size 826 bp. Two amplicons visible for all WT cells, no product seen for *TPSAN10* mutants. N= 3 technical repeats performed. **(H) RPE Directed differentiation protocol**.^21^ Subset of cells were grown on cover slips at passage 1, Day 35 and immunocytochemistry performed for early RPE markers. All differentiations of all clones showed early RPE markers. WT cells showed expression of TSPAN10 at Day 35 of directed differentiation. Top images are phase microscopy of cells at varying stages of differentiation. Scale bars 125 µm in microscopy images, scale bars 25 µm in immunocytochemistry images

In total nine independent differentiations of RPE from the three hESC cell lines (WT, A12 & A2) were generated, three WT, three A12 and three A2. Differentiations were staggered weekly, therefore one differentiation of WT, A12 and A2 was initiated over a period of three weeks. Directed differentiation was used to differentiate hESC into RPE, ^21^ as shown in Figure 1H. Directed differentiation was selected because TSPAN10 was hypothesised to influence RPE pigmentation, therefore impacting clone selection based on pigment levels.

Knockout of TSPAN10 in all differentiations of A2 and A12 was confirmed using immunocytochemistry (Figure 1 B - D), Western blotting (Figures 1 E & 1 F), and RT-PCR (Figure 1 G). WT cells showed expression of TSPAN10 during the differentiation process (Figure 1H) and continued to show expression once fully differentiated. In WT cells TSPAN10 was localised to the late endosome through colocalization with the late endosomal marker Rab7 (Supplemental Figure S5). To identify a phenotype caused by TSPAN10^-/-^ in RPE cells validation was first sought to confirm that hESCs were correctly differentiated to healthy RPE. All cells were therefore characterised for the presence of RPE morphological and functional features in vitro.

### Characterisation of hESC-RPE

All clones produced functional RPE monolayers on porous inserts with expression of RPE markers. All differentiations showed intact junctional localisation of occludin and zonula occludens (ZO-1) (Figure 2 A - F). There were no differences in the ability of TSPAN10^-/-^ and WT cells to phagocytose photoreceptor outer segments (POS), with equal numbers of POS attached and internalised in all clones and differentiations (Figure 3 G - I). TSPAN10^-/-^ cells appeared to be less well polarised and less columnar than their WT equivalents (Figure 3 J-R). All differentiations expressed the Na^+^/K^+^ ATPase (Figure 3 J – R, Supplemental Figure S5), however WT cells showed predominant apical expression, whereas expression in TSPAN10^-/-^ cells was more global evident of reduced polarisation. PEDF was secreted in a polarised manner, with increased apical: basal secretion in both TSPAN10^-/-^and WT cells. VEGF was predominantly basally secreted in WT cells; however, TSPAN10^-/-^ cells showed no significant differences between basal and apical secretion (Figure 3 M & N), likely reflecting reduced cell polarisation.

**Figure 2.**
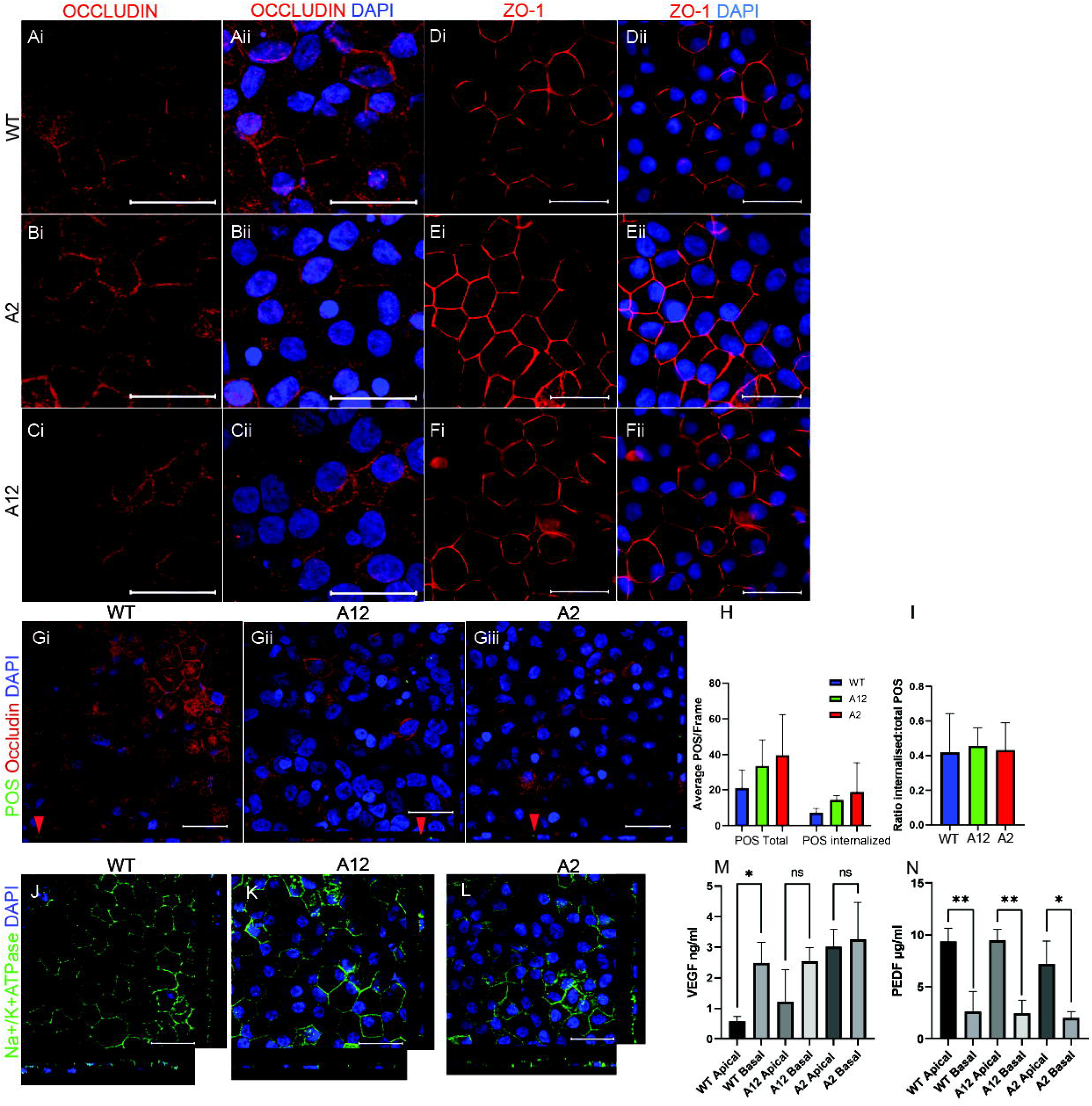
Phenotyping hESC-RPE for RPE characteristics A-C Immunocytochemistry for Occludin in hESC-RPE. Ai, Bi. Ci Representative staining of Occludin (red). Aii, Bii, Cii Merged images showing dual DAPI and Occludin staining. Scale bars represent 25µm Immunocytochemistry performed for three differentiations from all three clones (n=9). hESC-RPE cultured on transwells. **D - F Immunocytochemistry for ZO-1 in hESC-RPE** Di, Ei. Fi Representative staining of ZO-1 (red) Dii, Eii, Fii Merged images showing DAPI and ZO-1 staining Scale bars represent 25µm Immunocytochemistry performed for three differentiations from all three clones (n=9). hESC-RPE cultured on transwells. **G - I Phagocytic ability of hESC-RPE** G: Confocal z-stack images 4 hrs after feeding porcine POS labelled with FiTC. Red arrow heads indicate internalised POS (green). Occludin (red) and DAPI (blue) detected to visualise cells. Scale bar 25 μm Gi: WT, Gii: A12, Giii: A2. **H**: Bar chart showing quantification of average number of POS/frame, n = 3 cell lines with three replicate confocal frames (63× magnification) each. Error bars indicate SD. **I**: Bar chart showing ratio of internalised to total POS. n = 3 cell lines with three replicate confocal frames (63× magnification) each. Error bars indicate SD. **J - L: Immunocytochemistry for Na+/K+ ATPase in hESC-RPE** Confocal z-stack images of immunofluorescent labelling for the RPE marker Na^+^/K^+^-ATPase (green) and nuclear stain DAPI (blue) of hESC-RPE cultured on transwells (n=3 for each clone). J: WT, K: A12, L: A2 Predominantly apical localisation of Na^+^/K^+^-ATPase pump in WT cells. Apical localisation is lost in A2 and A12. Scale bars represent 25µm. **M & N Secretion of growth factors from hESC-RPE** **M: Secretion of vascular endothelial growth factor (VEGF) from hESC-RPE** ELISA quantification of VEGF was performed on conditioned media harvested from the apical and basal compartments of hESC-RPE cultured on transwell inserts. There was a statistically significant increase in basal vs apical conditioned media VEGF for WT cells. There were no statistically significant differences in basal vs apical conditioned media VEGF for the TSPAN10^-/-^ clones A12 and A2. Results are expressed as mean + SD for three separate differentiations for each cell clone. Statistical analysis performed using One way ANOVA with Tukey’s correction for multiple comparisons. \**p*<0.05 **N: Secretion of pigment derived epithelial growth factor (PEDF) from hESC-RPE** ELISA quantification of PEDF was performed on conditioned media harvested from the apical and basal compartments of hESC-RPE cultured on transwell inserts. Results are mean + SD for three separate differentiations for each cell clone. Each sample was run in duplicate. PEDF was significantly higher in apical compared to basal conditioned media for WT p=0.007; A12 p=0.0018; A2 p=0.018 Statistical analysis performed using One-way ANOVA with Tukey’s correction for multiple comparisons. **p<0.01, *p<0.05

**Figure 3.**
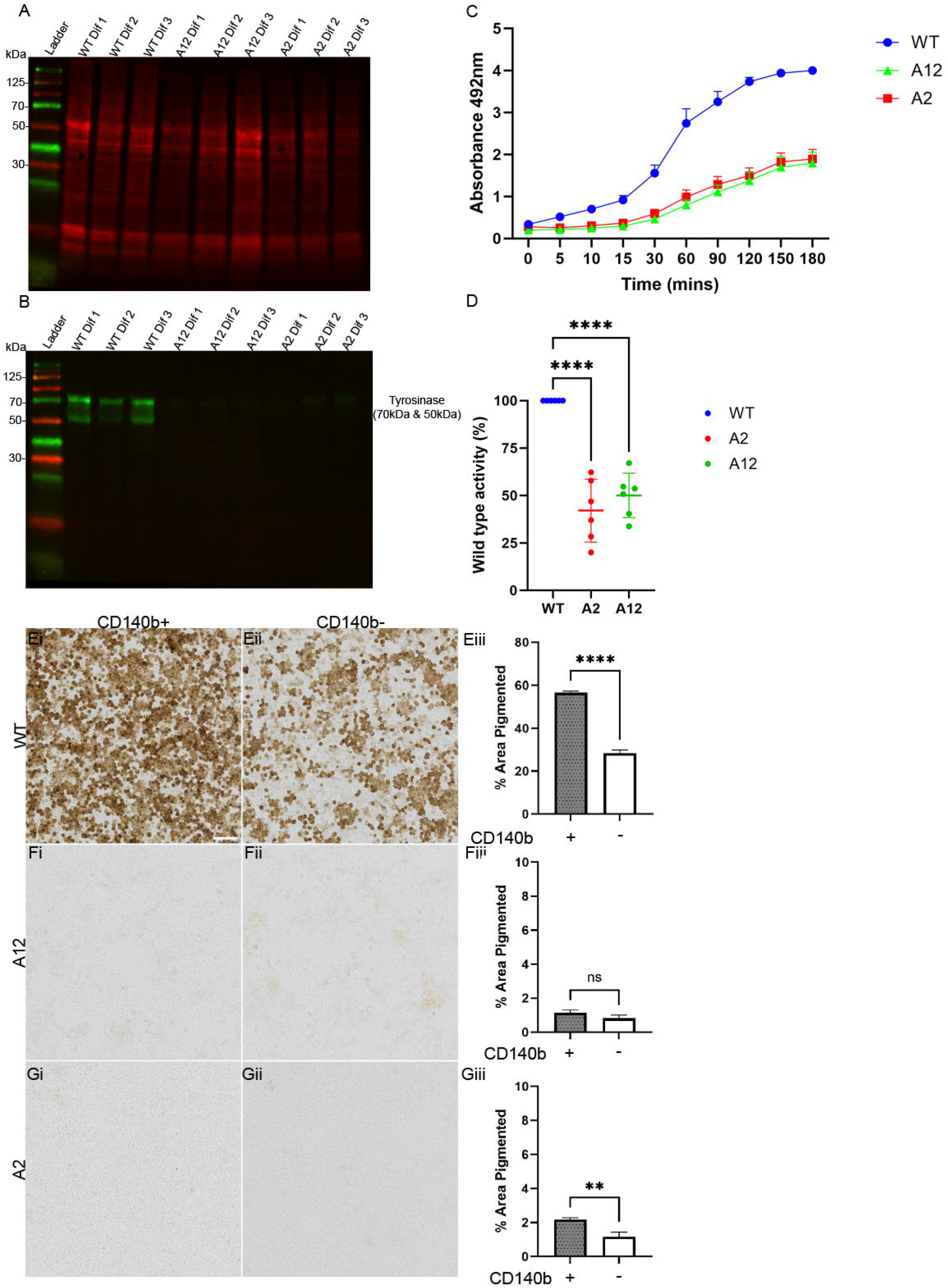
Study of pigmentation and tyrosinase function in hESC-RPE. **Western Blot for Tyrosinase in TPSAN10 mutant and WT hESC-RPE cells** **A:** A total protein stain (Revert 700) was performed as a control to ensure equal loading of lysates. **B:** A Tyrosinase protein blot (60kDa) indicates that tyrosinase protein expression is reduced in all differentiations of A12 and A2 in comparison to WT. N=3 technical repeats performed **Tyrosinase activity in wild-type and TSPAN10 mutant hESC-RPE** The absorbance of dopaquinone, a product synthesised by the transformation of L-DOPA by tyrosinase was quantified as a measure of tyrosinase activity in wild-type and TSPAN10-mutant cell lines (A2 and A12). Cumulative production of dopaquinone (**C**) was quantified from the start of L-DOPA treatment (0 min) to 180 min. Statistical differences between cell lines were analysed at 180 min (**D**). Data are shown as mean ± SD and statistically significant differences between groups are indicated by asterisks (****p < 0.0001). **FACS sorting for CD140b effects on cell pigmentation** **Ei** Brightfield image WT CD140b+ hESC-RPE **Eii** Brightfield image WT CD140b- hESC-RPE **Eiii** Bar chart depicts mean % area pigmented for WT CD140b + and – FACS sorted cells. Significant difference evident. **Fi** Brightfield image A12 CD140b+ hESC-RPE **Fii** Brightfield image A12 CD140b- hESC-RPE **Fiii** Bar chart depicts mean percentage (%) area pigmented for A12 CD140b + and – FACS sorted cells. No significant difference seen, cells have very little pigmentation. **Gi** Brightfield image A2 CD140b+ hESC-RPE **Gii** Brightfield image A2 CD140b- hESC-RPE **Giii** Bar chart depicts mean % area pigmented for A2 CD140b + and – FACS sorted cells. Significant difference seen although cells still have very little pigmentation. Scale bars represent 100µm Statistical analysis with Student’s two-tailed t-test ****p<0.0001, *** p<0.001, **p<0.005, *p<0.05

### TSPAN10^-/-^hESC-RPE show defects in pigmentation

A key characteristic of RPE cells is their dark brown/black pigmentation. TSPAN10^-/-^ RPE did not become pigmented during the differentiation process, whereas WT hESC-RPE formed classical polarised, pigmented cobblestone monolayers. Cells had previously been characterised for RPE markers including tyrosinase and no differences in *TYR* gene expression levels were seen (data not shown). Western blotting for tyrosinase however showed a significant reduction in TSPAN10^-/-^ cells in comparison to WT (Figure 3 A & B).

One of the most widely used in vitro assays to measure the dopa oxidase activity of tyrosinase is the dopachrome method ^22^. The formation of dopachrome from L-dopa is measured spectrophotometrically. TSPAN10^-/-^ cells showed a significant reduction in tyrosinase activity in comparison to WT throughout the assay, (Figure 3 C). By 180 minutes all WT differentiations had reached the maximum threshold at 490 nm absorbance, however both TSPAN10 mutants, A12 and A2 showed only approximately 50% of this activity at 180 minutes, Figure 3 D.

RPE cells sorted for CD140b expression have previously been reported to be more pigmented ^23^, therefore fluorescence activated cell sorting (FACS) of hESC-RPE was performed with sorted cells grown on RPE monolayers for high and low CD140b expression. WT hESC-RPE with high CD140b expression were significantly more pigmented than cells with low expression (Figure 3 E). However, no obvious differences in pigmentation were seen for either TSPAN10^-/-^ cell clone (Figure 3 F & 3 G). Further details of FACS parameters can be found in Supplemental Figure S7 and pigmentation thresholding in Supplemental Figure S8.

### Ultrastructural analysis of TSPAN10 k.o RPE reveals shorter cell bodies with reduced melanosome numbers

The ultrastructure of hESC-RPE monolayers cultured on permeable inserts was analysed by TEM. WT cells were found to be polarised in a typical apical–basal axis and showed characteristic RPE structures, including apical microvilli, melanosomes, basal nuclei and basolateral infolds, (Figure 4 A – C). TSPAN10^-/-^ RPE were less well polarised, as previously demonstrated in Figure 2, however all differentiations showed characteristic RPE structures, aside from differences in melanosomes, discussed below. There was a significant difference in cell body height between WT and TSPAN10^-/-^ RPE (A2 p<0.0001, A12 p<0.0001, One-Way ANOVA) (Figure 4 D), mutant cell bodies were shorter than their WT counterparts.

**Figure 4.**
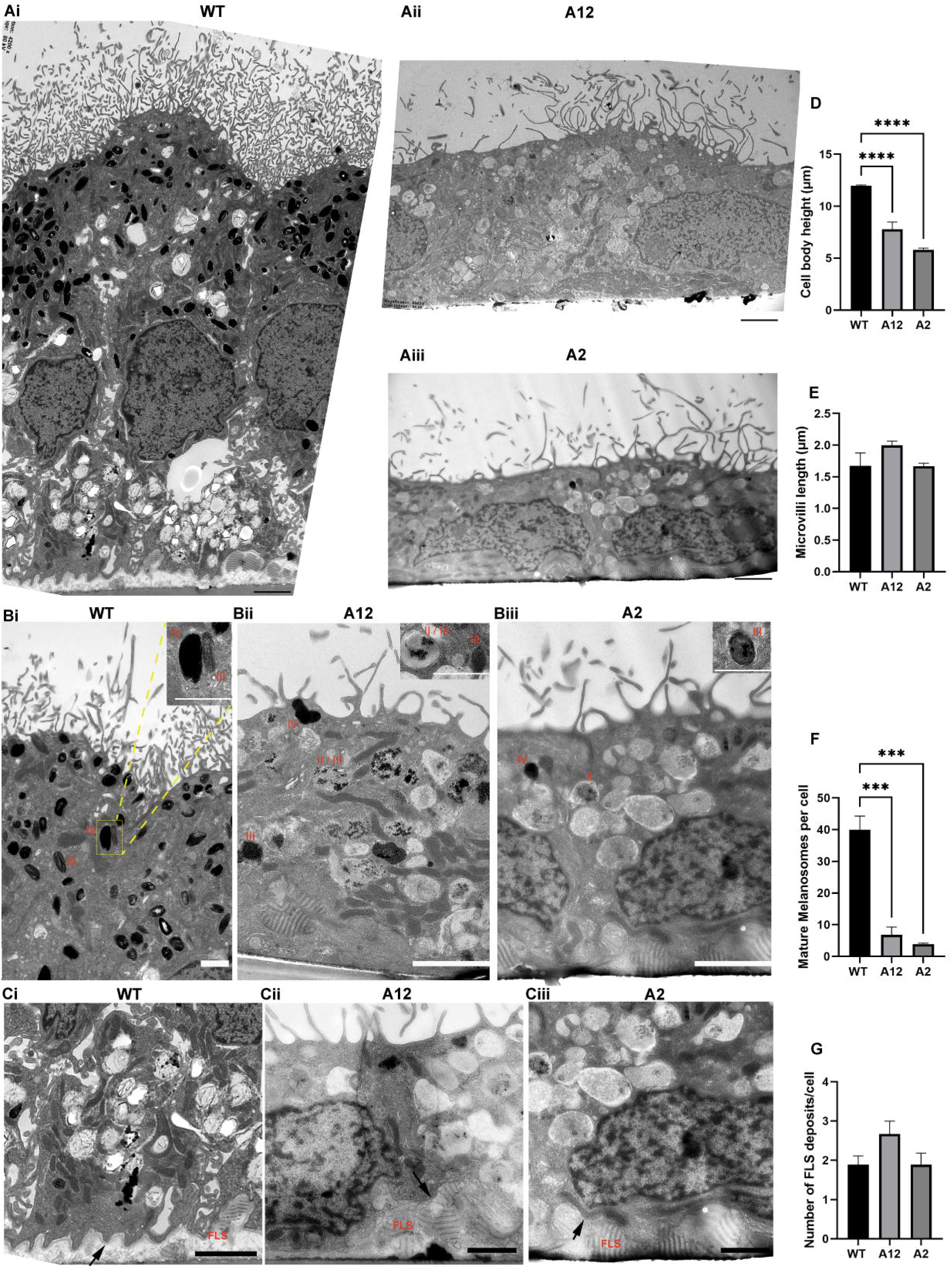
Ultrastructural Analysis of hESC-RPE. A significant difference was observed in cell body height between WT and TSPAN10 mutants (**Ai-Aiii**). WT cells displayed significantly increased numbers of stage III and IV melanosomes in comparison to TSPAN10^-/-^ cells (**Bi-Biii**), examples of melanosome staging shown from Stage II to Stage IV. There was no difference in the amount of sub-RPE fibrous-long spaced collagen (FLS) deposits between WT and TSPAN10^-/-^ cells, nor differences in basal laminar area (**Ci-Ciii**). Black arrows depict basal infolds. Abbreviations: FLS: Fibrous-long spaced collagen, II: Stage 2 melanosomes, III: Stage 3 melanosomes, IV: Stage 4 melanosomes. Scale bars in A represent 2 µm, B = 1 µm and in C, WT = 2 µm, A12 & A2 = 1 µm. Bar charts demonstrate average RPE characteristics identified using transmission electron microscopy (TEM). **D**: Significant reduction in RPE average cell body height in TSPAN10^-/-^ mutant clones in comparison to WT. **E**: Significant increase in the number of Stage III and IV (mature) melanosomes per cell for WT cells in comparison to TSPAN10 k.o mutants. **F**: No differences seen in the number of sub-RPE FLS collagen per cell **G**: No differences seen in the average apical microvilli length Three cells examined per image. Three differentiations per cell clone (N=9). For cell body height measurements three readings taken per cell and averaged. For apical microvilli measurements three microvilli measurements were taken per cell and averaged. Error bars represent SEM. Statistical analysis with One-way ANOVA **** p<0.001 ***p<0.005

Furthermore, there was a significant reduction in the number of mature melanosomes between TSPAN10^-/-^ and WT hESC-RPE (Figure 4 B & E) (A2 p=0.0002, A12 p=0.0004, One-Way ANOVA), with a reduction in ovoid melanosome shape seen in TSPAN10 mutant cells. No differences were seen in the average length of apical microvilli (p = 0.1996) (Figure 4 A & G). Sub-RPE deposits with a distinct striated pattern were identified as fibrous long-spacing (FLS) collagen and were observed between RPE and the underlying insert surface in all samples (Figure 4 C, arrows). No difference in the number of FLS collagen deposits per cell was observed between TSPAN10^-/-^ and control-hESC-RPE (p = 0.1168) (Figure 4 F).

### RNA sequencing analysis of TSPAN10^-/-^ RPE in comparison to WT

RNA-seq was used to investigate gene expression in the TSPAN10 mutants A12 and A2 in comparison to WT hESC-RPE. The aim was to identify transcriptional changes common to A12 and A2 that occurred because of TSPAN10^-/-^.

Log2 fold-change data were subjected to two analysis pipelines: (i) differential gene expression analysis, to investigate individual genes of interest, and (ii) gene set enrichment analysis (GSEA), to identify over-represented sets of genes.

Genes with an absolute log2 fold-change >1.00 and a padj <0.05 were called as differentially expressed genes (DEGs). A12 had 489 DEGs in comparison to WT, were as A2 had 152.

Augmented differential gene expression was not surprising in both A12 and A2 given the previously described phenotypes. Compared to WT, A12 and A2 shared 49 upregulated and 64 downregulated DEGs, representing changes assumed to be caused by the loss of TSPAN10. A heatmap showing the top 50 key up and down-regulated genes is shown in Figure 5.

**Figure 5.**
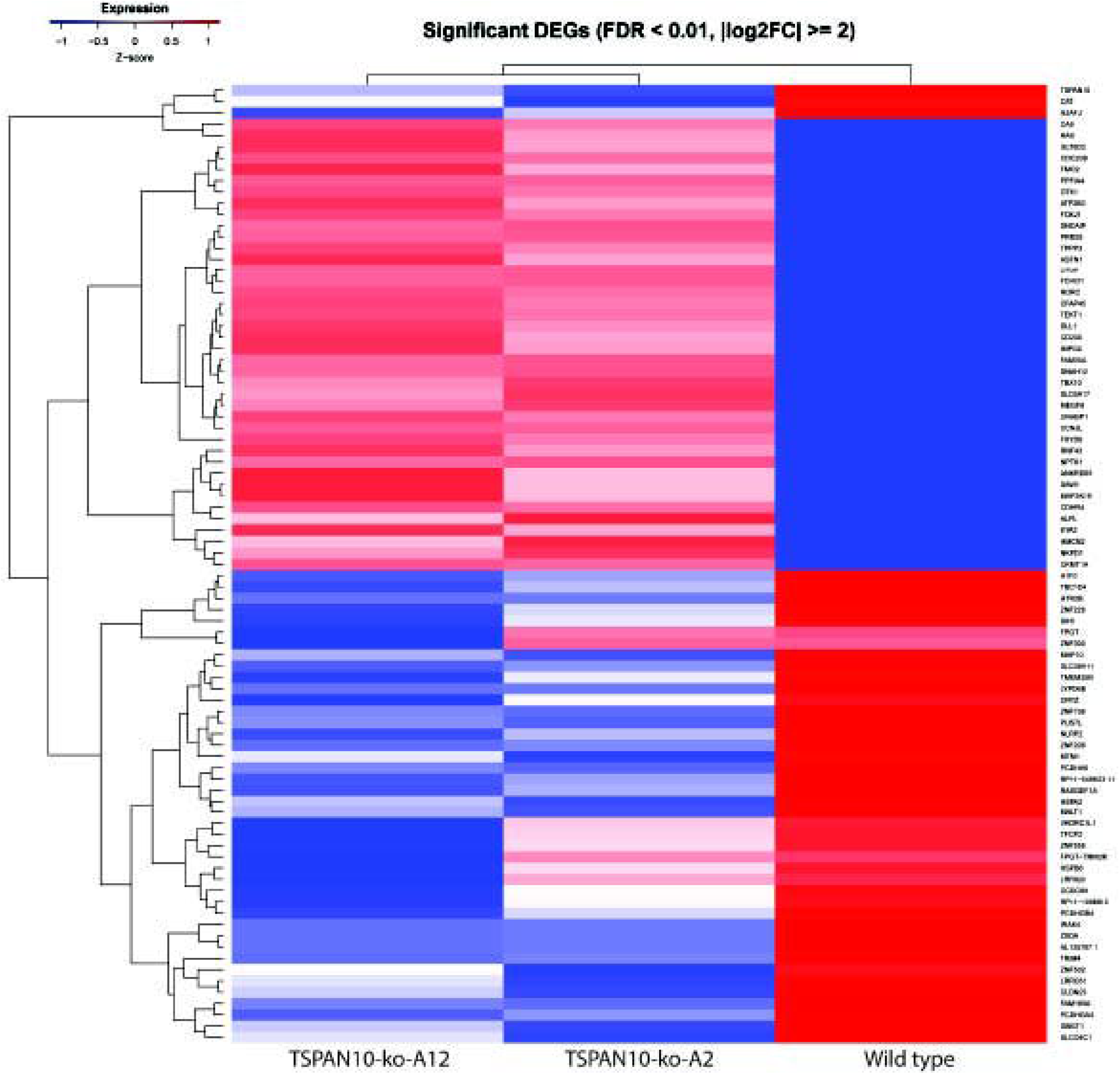
Transcriptomic profiling reveals gene expression changes in TSPAN10^-/-^ RPE. Heatmap demonstrating hierarchical clustering of the top 50 significant differentially expressed genes (DEGs) between two independent TSPAN10^-/-^ RPE cell lines (A2 and A12) and a WT control. Gene expression levels were obtained via RNA sequencing of hESC-RPE. Significance was defined using strict thresholds: False Discovery Rate (FDR) < 0.01 and an absolute log2 fold change (|log2FC|) ≥ 2. Colour scaling represents the Z- score of normalized read counts, ranging from blue (lower expression, Z-score ≤ −1) to red (higher expression, Z- score ≥ 1). Both column (sample) and row (gene) dendrograms illustrate clustering based on expression similarity. Genes of interest include *TSPAN10, CAT* and *HSPA2*, which were all significantly downregulated in knockout cells.

While transcriptomic profiling revealed low-level expression of a truncated *TSPAN10* transcript in knockout lines (Figure 5), we detected no functional protein product (Figure 1), an observation consistent with previous reports of residual non-functional transcripts in similar CRISPR/Cas9 mediated knockout models ^24^.

Those genes with the greatest fold change for both A12 and A2 versus WT were examined. Catalase (*CAT*) was significantly down-regulated in both A2 (log2 fold-change −2.33, p_adj_ = 1.39 x 10^-20^) and A12 (log2 fold-change −1.52, p_adj_ = 1.05 x 10^-7^) compared to WT. Catalase is a key enzyme in the cell which catalyses the decomposition of hydrogen peroxide ^25^.

Catalase activity has previously been reported to be associated with AMD ^25^. GO pathway enrichment showed terms related to misfolded protein binding to be negatively enriched in both TSPAN10^-/-^ clones. Within this gene set DEGs included several heat shock proteins, namely HSPA2 (A2 log2fold= −3.55, padj=0.03, A12 log2fold= −2.89, padj=0.02), HSPA1A (A2 log2fold= −1.03, padj=0.01, A12 log2fold= −1.55, padj=1.22 x 10^-5^) and HSPA13 (A2 log2fold = −1.39, padj=0.007, A12 log2fold = −1.12 padj=0.006). Both catalase and heat shock proteins are involved in the oxidative stress response of the cell, consequently we examined this in our TSPAN10^-/-^ and WT hESC-RPE.

The full RNAseq dataset has been deposited in NCBI’s Gene Expression Omnibus^26^ and is accessible through GEO Series accession number GSE352284. (https://www.ncbi.nlm.nih.gov/geo/query/acc.cgi?acc= GSE352284).

### Oxidative stress response changes in TSPAN10^-/-^ RPE in comparison to WT

Western blotting for catalase confirmed our RNAseq data with reduced catalase expression seen in TSPAN10^-/-^ versus WT hESC-RPE (Figure 6 A & B). The enzymatic activity of catalase was studied in hESC-RPE in basal conditions. A significant reduction in the activity of catalase was observed in TSPAN10^-/-^ cells in comparison to WT (Figure 6 C).

**Figure 6.**
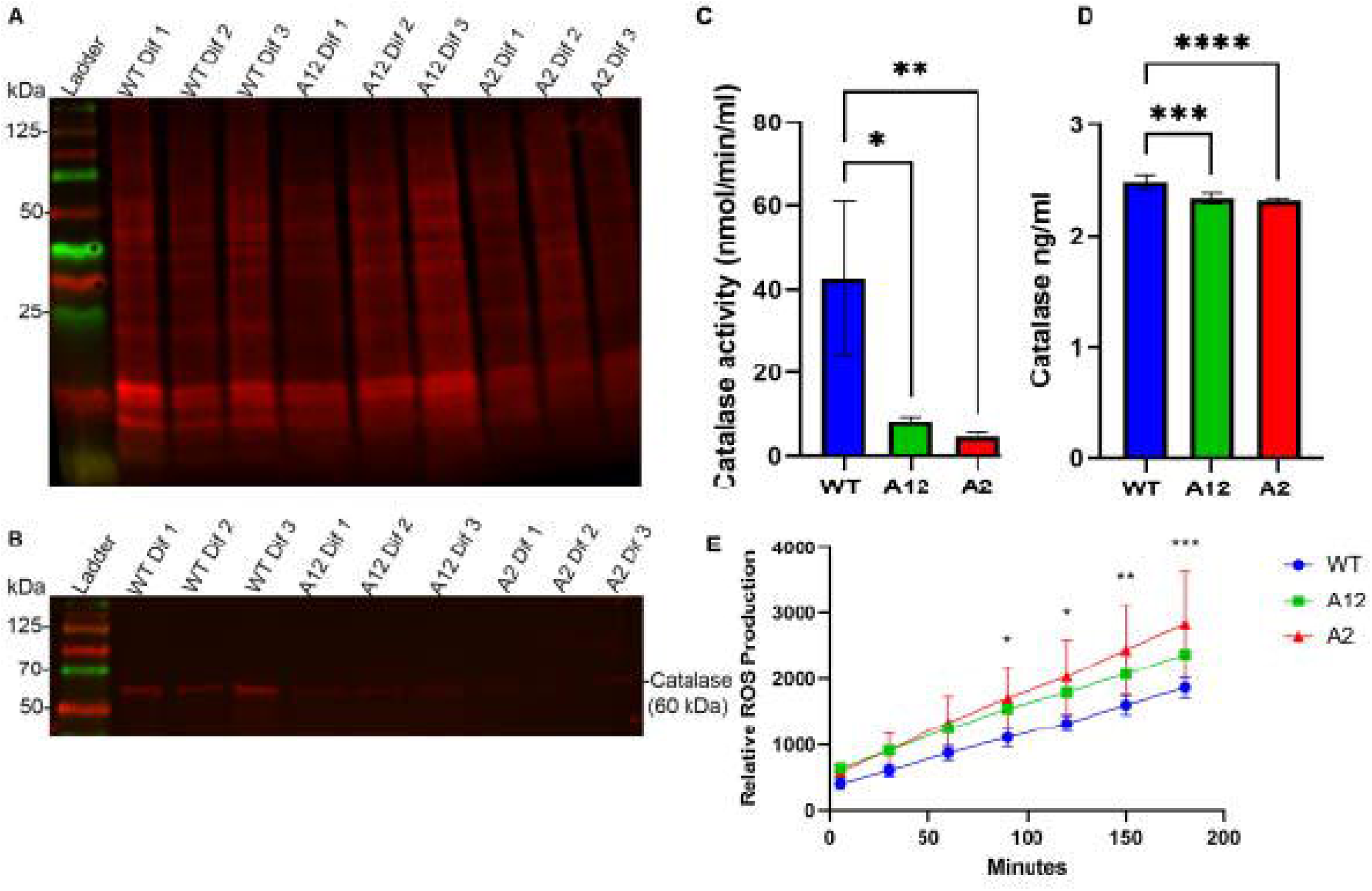
Study of oxidative stress in hESC-RPE. Western Blot for Catalase the peroxisome marker in TPSAN10 mutant and WT hESC-RPE cells **A:** A total protein stain (Revert 700) was performed as a control to ensure equal loading of lysates. **B:** A Catalase protein blot using the Abcam monoclonal Ab indicates that catalase protein expression is reduced in all differentiations of A12 and A2 in comparison to WT. N=3 technical repeats performed. **C: Determination of the antioxidant enzyme catalase activity in hESC-RPE** Catalase activity during basal conditions determined for wild-type and the TSPAN10 knockout cell lines A12 and A2. Enzyme activity expressed as means ± SD of three differentiations for each cell line, where each differentiation was repeated in triplicate. Statistical significance with respect to WT cells using One-way ANOVA with Dunnett’s multiple comparison post-hoc test. p ≤ 0.05 (*), p ≤ 0.01 (**) **D**: ELISA quantification of catalase was performed on conditioned media harvested from the apical compartments of hESC-RPE cultured on transwell inserts. Bars show mean of three differentiations for each cell clone. Error bars show SD of three differentiations. One-way ANOVA with Dunnett’s multiple comparison post- hoc test used for statistical analysis. ****p<0.0001, ***p=0.0002, *p<0.05 **E: TSPAN10 knockout hESC-RPE produce increased reactive oxygen species (ROS)** DCFDA assay in WT, A12 and A2. Reactive oxygen species (ROS) production is significantly higher in TSPAN10 knockout hESC-RPE under basal conditions in comparison to WT. Statistical comparison conducted using a two-way ANOVA with Dunnett’s multiple comparison test. Significant differences are indicated as p ≤ 0.05 (*), p ≤ 0.005 (**) and p ≤ 0.001 (***). Abbreviations: DCFDA

Furthermore, ELISA for catalase in conditioned media from the apical compartments of hESC-RPE showed a statistically significant reduction in TSPAN10^-/-^ cells (Figure 6 D). Given that catalase is a key antioxidant enzyme in the cell we compared reactive oxygen species (ROS) production in WT and TSPAN10^-/-^ cells. Figure 6 E demonstrates increased ROS production at baseline in TSPAN10^-/-^ cells in comparison to WT over 180 minutes.

Taken together these results emphasise the reduced antioxidant enzyme production and consequent increased oxidative stress present in TSPAN10^-/-^ cells in comparison to WT.

### Summary of TSPAN10^-/-^ Phenotype

TSPAN10^-/-^ cells were differentiated to mature RPE, expressing key RPE markers and able to perform known RPE functions. However, there were several phenotypic differences to WT cells. To summarise lack of TSPAN10 results in: loss of pigmentation, an increase in oxidative stress, reduced polarisation, and ultrastructural changes resulting in shorter RPE cells with reduced melanocytes (Supplementary Figure S-9).

## Discussion

The molecular pathophysiology of AMD remains poorly understood. AMD is a complex disease with both genetic and environmental contributors. The specific role of many AMD risk genes, such as *TSPAN10*, is not currently known. Here we investigated TSPAN10 in human stem cell derived RPE utilising CRISPR/Cas9 gene editing.

TSPAN10, a member of the tetraspanin family, has been increasingly recognised for its involvement in cellular processes such as membrane organisation, signal transduction, and cell morphology ^27^. Our study provides novel insights into the role of TSPAN10 in the RPE, a critical layer responsible for supporting photoreceptor function and maintaining retinal homeostasis.

One of the most striking observations was the severe hypopigmentation in TSPAN10^⁻/⁻^ RPE cells, which occurred across all differentiations, independent of CD140b expression. This was substantiated at the molecular, functional, and ultrastructural levels. SNPs in *TSPAN10* have been associated with lighter hair and eye colour ^15–18^, and recently a significant association was seen between TSPAN10 and the amount of fundus pigmentation ^19^, aligning with our results. By knocking out TSPAN10 in hESC-RPE we saw a significant reduction or even absence of RPE pigmentation. Although *TYR* mRNA levels were unaltered, tyrosinase protein expression and enzymatic activity were both markedly reduced.. While preliminary immunocytochemistry in WT cells showed some spatial overlap between TSPAN10 and Rab7, a marker of late endosomes, the exact role of TSPAN10 within the endosome remains to be definitively understood, including how it affects melanogenic enzymes such as tyrosinase and its role in melanosome biogenesis ^28^. Nonetheless, the marked reduction in mature melanosomes and total absence of pigmentation in knockout cells mirrors phenotypes seen in pigmentation-related disorders involving trafficking defects ^29,30^..

Epidemiological data indicates a higher prevalence of AMD in populations with reduced ocular pigmentation ^31–34^. While our in vitro model demonstrates that TSPAN10 deficiency results in severe RPE hypopigmentation, determining whether TSPAN10 variants contribute to demographic disparities in AMD incidence requires further in vivo investigation..

Interestingly, FACS using CD140b, a recently identified marker correlated with RPE pigmentation^23^, revealed that while the proportion of CD140b-positive cells was comparable between TSPAN10^⁻/⁻^and wild type populations, pigmentation levels within the CD140b positive subpopulation were still significantly diminished in the knockout cells. However, WT cells showed an obvious difference in pigmentation levels between CD140b positive and negative populations. This dissociation suggests that TSPAN10 is not essential for initial RPE lineage specification or the expression of pigmentation-associated surface markers but is critical for the maturation and functional pigmentation capacity of these cells.

TSPAN10^-/-^ RPE showed characteristics of mature RPE with expression of key RPE markers. Importantly, although tight junction proteins ZO-1 and occludin were expressed in both WT and knockout cells, Na^+^/K^+^ ATPase a marker of cell polarity was evident predominantly apically in WT cells, however in TSPAN10^-/-^ cells this polarisation was lost. Furthermore, the loss of polarisation was functionally evident, most notably through the non- polarised VEGF secretion observed in TSPAN10^⁻/⁻^ RPE. This contrasts with the basal VEGF bias in WT RPE and may have implications for the association of TSPAN10 in AMD, specifically nAMD. Conversely, PEDF secretion remained apically polarized in both WT and TSPAN10^-/-^ cells. The preserved phagocytosis of photoreceptor outer segments in TSPAN10-deficient RPE suggests that not all core functions are impaired, indicating a selective, rather than global, influence on RPE functionality.

Ultrastructural analysis further revealed morphological alterations in TSPAN10^⁻/⁻^ RPE, including shorter cell bodies, and fewer mature melanosomes, with no ovoid melanosomes seen. These ultrastructural alterations imply that TSPAN10 may contribute to melanosome biogenesis or trafficking pathways. Furthermore, the reduction in cell body height seen corroborates our clinical study ^6^, which correlated the presence of the AMD risk SNP at TSPAN10, rs6565597 with reduced thickness of the RPE-BM segment on optical coherence tomography, reinforcing the relevance of our model to human disease. When interpreting our ultrastructural findings, it is important to consider the morphological differences between native RPE and in vitro models. Recent volume electron microscopy studies have demonstrated that in native human tissue, mature spindle-shaped melanosomes are predominantly localised within the elongated apical processes, contributing significantly to the interdigitation zone band observed on clinical OCT. In contrast, our hESC-RPE model, consistent with many cultured RPE systems, exhibits mature melanosomes that are primarily constrained within the main cell body, accompanied by more stubby microvilli.

Transcriptomic profiling of TSPAN10^⁻/⁻^ RPE identified consistent downregulation of genes involved in protein folding and oxidative stress responses, including several heat shock proteins and catalase. These findings were validated at the protein and functional levels: catalase expression and activity were significantly reduced in the knockout lines, and this was accompanied by elevated basal ROS levels, indicating a shift toward oxidative stress. This is particularly relevant given the RPE’s high metabolic demand and susceptibility to oxidative damage over a lifetime. Loss of TSPAN10 may compromise intrinsic antioxidant defences or disrupt intracellular signalling cascades involved in redox homeostasis, leading to increased cellular vulnerability. Given that oxidative stress is a key driver of RPE degeneration in aging and AMD, our findings offer a compelling mechanistic link between reduced TSPAN10 expression, redox imbalance, and AMD susceptibility.

The link between pigmentation and oxidative stress adds an additional layer of complexity. Melanin serves as a potent antioxidant ^35^, with RPE melanin known to play a central cytoprotective role as a scavenger of oxidative stress in the retina ^36^. The hypopigmentation observed in TSPAN10-deficient cells stemming from reduced melanosome numbers and melanin may thus exacerbate oxidative stress. Conversely, elevated ROS levels may impair pigmentation processes. Reduced catalase activity, an enzyme known to modulate tyrosinase function via H₂O₂ detoxification, may further link oxidative stress with impaired melanogenesis ^37^. The possibility that TSPAN10 acts upstream of catalase or other stress- response elements raises intriguing questions regarding its broader regulatory functions.

Taken together, our data suggest that TSPAN10 is essential for multiple RPE-specific functions: (i) pigmentation and melanosome maturation, likely through late endosomal trafficking of tyrosinase; (ii) maintenance of epithelial polarisation, with implications for directional secretion of growth factors; and (iii) oxidative stress, through regulation of antioxidant enzymes such as catalase. Importantly, genetic variants in TSPAN10 have been associated with AMD, and our results provide experimental support for a causal role of TSPAN10 in RPE pathology. This establishes a valuable in vitro model for AMD-related dysfunction and suggests that modulating TSPAN10 activity or its downstream pathways could be a promising therapeutic avenue.

Future work should aim to delineate the precise trafficking role of TSPAN10 in the RPE, explore whether its loss sensitises RPE to environmental insults, and evaluate whether overexpression or rescue of TSPAN10 can restore RPE function in knockout models.

Additionally, given the complex regulation of oxidative stress in the retina, strategies to boost catalase activity or reduce ROS may mitigate the cellular stress observed in TSPAN10- deficient cells.

## Methods

### Culture of Human Embryonic Stem Cells

The MasterShef7 human embryonic stem cell line (passage 8), generated by the Centre for Stem Cell Biology, University of Sheffield, was cultured using standard feeder cell-free conditions, on Laminin-521 (LN521) (Biolamina) coated T25 flasks (maintenance) or 6-well plates (electroporation, differentiation). Flasks were coated with 5 µg/mL LN521 in phosphate-buffered saline (PBS) (+Mg/+Ca^2+^, PBS Life Technologies, #14190-091) at 4°C overnight or 2 hours at 37°C. Laminin solution was aspirated and the flask covered with the pluripotent stem cell medium NutriStem XF (Biological Industries, #05-100-1A). Cells were seeded at 10000 cells/cm^2^ and left for 48 hours and then fed daily. The cells were passaged every 3-5 days at approximately 80% confluency. When passaged the cells were washed twice with PBS (-Mg/-Ca^2+^, PBS Life Technologies #14190-094), and detached by incubating with a 0.5 mM solution of EDTA (Life Technologies, #15575-020) in PBS^-/-^ for 5 minutes at 37°C. After addition of 5 mL Nutristem XF, the cell suspension was centrifuged for 5 minutes at 200 g and counted before seeding into a fresh flask.

### CRISPR/Cas9 Gene Editing

The Benchling CRISPR finder tool was used with default setting to identify sgRNA sequences within 10 bp of the TSPAN10 ATG start signal in exon 2. The TSPAN10 sgRNA sequence (GTGGCTGTCTGATAAGCTGA) was cloned under the control of a U6 promoter into a modified version of the chimeric CRISPR plasmid pSpCas9(BB)-2A-Puro (PX459) V2.0 (Addgene, # 62988), in which the ChB promoter was replaced with an EF1α promoter, to enhance expression in human pluripotent stem cells. 1 × 10^6^ cells (passage 18) in 100 µl Optimem medium were co-electroporated with the Cas9 expression plasmid containing the guide (5 µg) using a BTX ECM 830 electroporator (100V, 5 ms, 50 ms interval, dual pulse). Twenty-four hours after electroporation, the cells were subjected to 0.129 ug/ml puromycin (INVIVOGEN) selection for 72 hours until only single-cell clones remained. After a recovery phase of 1 week, clearly demarcated clones were picked and split individually into two 96- well plates to produce replicates that were then used for either genotyping or clonal expansion. gDNA was extracted using QuickExtract (Lucigen, #QE09050). PCR was performed using spanning primers as well as 5 and 3 primers for genome/construct junctions, which are listed in the supplemental methods, Table S-2. Two independent clones (A2, A12) with the desired mutation were identified out of 48 clones screened. Array comparative genome hybridization (array-CGH) was performed using Oxford Gene Technology’s (OGT) CytoSure Constitutional v3 60K oligo array, according to the manufacturer’s instructions, using Kreatech’s pooled control DNA as a reference (Kreatech Diagnostics, Amsterdam, Holland). Slides were scanned using a G2539A Agilent microarray scanner (Agilent Technologies, Wokingham, UK) and analyzed using OGT’s CytoSure Interpret (v4.9.40) microarray software.

### Differentiation of RPE

A directed differentiation approach was selected ^21,38^ and followed. Further details of the timeline and various steps in the directed differentiation protocol can be found in supplemental methods and a timeline depicting various passaging steps is shown in Figure 2 – 2. At day 84 cells had reached the end of the differentiation protocol ^19^ and were therefore considered to be fully differentiated into RPE and ready for cryopreservation and/or use in experiments.

For further analysis, hESC-RPE cells were cultivated for a minimum of 60 days on 12-well or 6-well transwell filter inserts (0.4-µm pore size, Corning Costar by Sigma-Aldrich, Munich, Germany) coated with GFR-Matrigel at a dilution of 1:30. hESC-RPE clones (n = 3 control; n = 3 A12; n = 3 A2) were characterised for the expression of cell-specific markers, transepithelial electrical resistance (TEER), polarised secretion of pigment epithelium-derived factor (PEDF) and vascular endothelial growth factor (VEGF) and phagocytic capacity.

### Transmission electron microscopy

For transmission electron microscopy (TEM) analysis, hESC-RPE (n = 3 control; n = 3 A12; n = 3 A2) were cultured on Transwell® inserts for 60 – 70 days. Details of sample preparation have been described previously ^39^.

### Western blotting

Western blotting was used to investigate the expression of TSPAN10 (1:100, 14430-1- AP;ProteinTech, Manchester, UK), tyrosinase (1:250, ab738; AbCam, Cambridge, UK), catalase (1:1000, ab76024; AbCam, Cambridge, UK) and total protein (LI-COR Biosciences 2018.REVERT™ Total Protein Stain) in hESC-RPE (n = 3 control; n = 3 A12; n = 3 A2, 60– 70 days) under reducing conditions and visualised with a LiCOR Odyssey imaging system.

### RT-PCR

Total RNA was extracted using the RNeasy Micro Kit (Qiagen) with on-column DNase digestion. cDNA was synthesised using the RT-NanoScript2 kit (Primerdesign). PCR amplification was performed using Bioline MyFi Mix (SLS Biosciences) and TSPAN10 primers 5’-ACAGAGGAGCCAGCGAAC-3’ and 5’-TGACAGAGGCTCCATCTTCG-3 with standard cycling conditions (35–40 cycles) and optimised annealing temperatures. Products were resolved on 1% agarose gels containing SybrSafe (Fisher Scientific) and imaged using a LiCOR Odyssey system.

### ELISA

Conditioned media were harvested from the apical and basal compartments of Transwell®- cultured hESC-RPE 60 h post-medium change. Secreted factors were quantified using commercial ELISA kits according to manufacturers’ instructions: Human PEDF (BioVendor RD191114200R; 1:8000 dilution), Human Catalase (Abcam ab277396; 1:25 dilution), and Human VEGF (Novex®; 1:10 dilution). Absorbance was measured at 450 nm with 570 nm wavelength correction using a SPECTROstar Nano microplate reader.

### Catalase Enzymatic Activity Assay

Catalase activity was quantified in hESC-RPE cell lysates using a Catalase Assay Kit (Cayman Chemical) according to the manufacturer’s instructions.

### Immunocytochemistry

hESC-RPE cells cultured on Transwell® inserts and hESC cells cultured on sterile glass cover slips were fixed in 4% PFA and blocked/permeabilized in PBS containing 0.2% Triton X-100, 5% BSA, and 1% donkey serum. Cells were incubated overnight at 4°C with primary antibodies targeting TSPAN10 (1:100, 14430-1-AP; ProteinTech, Manchester, UK), Rab7 (1:50, ab50533; AbCam, Cambridge, UK), ZO-1 (1:1000, 21773-1-AP; ProteinTech, Manchester, UK), Occludin (1:100, 71-1500; ProteinTech, Manchester, UK), Na^+^/K^+^ ATPase (1:100, ab7671; AbCam, Cambridge, UK), hNANOG (1:100, AF1997; R&D Systems, Minneapolis, MN, USA) SOX-2 (1:100, AF2018, R&D Systems, Minneapolis, MN, USA). Samples were labelled with species-specific AlexaFluor-conjugated secondary antibodies, counterstained with DAPI (Sigma Aldrich, UK), and mounted in Mowiol 4-88 (Sigma Aldrich, UK).

### Confocal Imaging

A Leica TCS-SP8 microscope with a 63x glycerol objective was used. Z-stacks were acquired using sequential scanning and system-optimised spacing. For co-localisation assays, co-localisation was analysed using Volocity software (PerkinElmer) as previously detailed ^40^. After background subtraction, automatic thresholding was applied using the Costes et al. algorithm ^41^ to calculate Manders’ split coefficients (M1 and M2).

### Phagocytosis Assay

FITC-labelled photoreceptor outer segments (POS) were isolated from fresh porcine retinas and quantified as previously described ^40^. Phagocytosis was assessed in mature hESC-RPE monolayers using a synchronised pulse-chase protocol adapted from previous work ^40^.

Briefly, cells were chilled and challenged with FITC-POS (4µg/cm^2^) at 17°C for 30 min to synchronise binding. Unbound POS were removed, and cells were incubated with prewarmed X-VIVO 10 at 37°C for 4h. Immunostaining for Occludin was performed and imaged using a Leica SP8 confocal microscope as described ^39^. Total and internalised POS were quantified using Fiji (version 2.9.0)^42^.

### DFCDA Assay

Intracellular ROS generation was determined using the 2’-7’dichlorofluorescein diacetate (DCFH-DA) assay (Abcam). Mature hESC-RPE cells were cultured in Nunc 96-well black optical-bottom plates (ThermoFisher). The assay was performed as per Manufacturer’s guidance; briefly, cells were incubated with 10 µM DCFH-DA for 40 min at 37°C and washed twice with PBS. Fluorescence intensity was measured using a microplate reader at 30- minute intervals up to 180 minutes.

### DOPA-Oxidase Activity Assay

DOPA-oxidase activity was assessed in all differentiations of all cell clones. Briefly cells were lysed using 1X radioimmunoprecipitation assay (RIPA) buffer (Sigma-Aldrich) + 1X cOmplete protease and phosphatase inhibitor cocktail (Roche), and the protein concentration was measured by BCA assay (Thermo Scientific™ Pierce™ BCA Protein Assay Kit). Samples were then diluted to 4 µg/µl and 30 µl used for DOPA-oxidase assays.

Samples were added to a 96 well plate in duplicate and 150 µl of a phosphate buffer (Sigma) with l-3,4-dihydroxyphenylalanine L-DOPA 1 mM (Sigma) and 3-methyl- 2- benzothiazolinonehydrazone hydrochloride 6 mM (MBTH) (Sigma) added. DOPA-oxidase activity was then measured as the accumulation of the downstream product, dopachrome.

The enzymatic activity was recorded by measuring the absorbance of dopachrome at 492 nm using a plate reader (SPECTROStar Nano; BMG Labtech, UK). This was performed at regular intervals at 37°C from the start of the L-DOPA treatment at 0 min for 30 min intervals until 180 min.

### Slide Scanner Microscopy

An Olympus BX51 microscope with robotic slide loader was used to capture brightfield images of slides. Initially an overview scan was taken. Five 1 x 1 mm squares were then selected for imaging transwell sections in detail. Three of the 1 x 1 mm squares were selected for each slide for analysis, any section with image distortion or blurring was not included. Three slides were imaged for each cell population.

### Pigmentation Image Analysis

Brightfield images were anonymised and analysed using Fiji (version 2.9.0)^42^. Initially to measure the level of pigmentation the background pixel values were set to 0 and black. Images were then duplicated, and the colour channels split, with the red channel selected. The black dots, representing transwell pores were thresholded, with maximum threshold based on the least pigmented image and set to 90. Thresholded selections were then transferred to the original image. The original image was converted to an 8-bit grey image to allow thresholding and calculation of the mean grey value, representative of pigment. A threshold of 0 – 205 was selected based on the least pigmented image. The threshold was then applied to create a binary image. This process was repeated for each image and the mean grey values calculated ± SD.

### Flow Cytometry and Fluorescence Activated Cell Sorting (FACS)

hESC-RPE monolayers were dissociated into single cells using a Papain Dissociation System (Lorne Laboratories). Cells were blocked in 1% BSA (Sigma) for 30 min on ice and immunostained with PE-conjugated anti-human CD140b (20 µg/mL; clone 28D4, BD Biosciences) and Fixable Viability Dye eFluor® 506 (0.5 µL/sample; Invitrogen) for 30 min at 4°C. Following two washes, cells were resuspended in 1% BSA and sorted on a BD FACS Aria Fusion sorter analysed with BD FACSDiva software (BD Biosciences). Spectral overlap was corrected using compensation beads.

### RNA Sequencing and Analysis

Total RNA was extracted from hESC-RPE cell pellets using the RNeasy Micro Kit (Qiagen) following the manufacturer’s protocol.

Library preparation and sequencing were performed by Novogene. mRNA was isolated using poly-A enrichment, randomly fragmented, and converted to cDNA using random hexamers. Following end-repair, A-tailing, and adapter ligation, libraries were sequenced on an Illumina NovaSeq 6000 platform using a paired-end 150bp (PE150) strategy.

Differential expression analysis was performed using the DESeq2 R package (v1.20.0). P- values were adjusted using the Benjamini-Hochberg method to control the false discovery rate (FDR), with an adjusted p ≤0.05 threshold for significance. Gene Ontology (GO) and Kyoto Encyclopedia of Genes and Genomes (KEGG) pathway enrichment analyses were implemented using the clusterProfiler R package. Functional enrichment was further assessed using Gene Set Enrichment Analysis (GSEA).

### Statistics

Statistical analyses were performed using GraphPad Prism (version 9.0). All data are presented as the mean ± standard deviation (SD) or mean ± standard error of the mean (SEM), as indicated in the respective figure legends. For all experiments, a minimum of three independent differentiations (n = 3) were performed for each cell line (WT, A12, and A2) to ensure biological reproducibility. Quantitative data comparing the three groups (WT vs. TSPAN10^-/-^ clones) were analysed using a one-way ANOVA followed by Tukey’s or Dunnett’s post-hoc tests for multiple comparisons. Specific comparisons between two groups were conducted using a Student’s two-tailed t-test. For time-course data, such as the reactive oxygen species (ROS) production assay, a two-way ANOVA with Dunnett’s multiple comparison test was applied. RNA-sequencing data were analysed using two pipelines: gene set enrichment analysis (GSEA) and differential gene expression analysis, with differentially expressed genes (DEGs) defined by an absolute log2 fold-change > 1.00 and an adjusted p-value (padj) < 0.05.

## Supplemental Information

Figures S1–S10, and Table S1.

## Resource Availability

Lead contact: Professor Andrew Lotery. This study generated two TSPAN10 knockout hESC lines These lines are available from the Lead Contact upon request with a complete Materials Transfer Agreement (MTA) dependent on stock availability.

## Acknowledgements

This study was supported by the Wellcome Trust [210572/Z/18/Z] and the Gift of Sight Charity. The views expressed are those of the author(s) and not necessarily those of the funders. We thank the Centre for Stem Cell Biology, The University of Sheffield, Sheffield, UK for ESC lines.

## Author Contributions

Rebecca Kaye: Conceptualisation, Methodology, Data Curation, Formal Analysis, Writing- original draft, Accountable for all aspects of work Liliya Nazlamova: Methodology, Data Curation Eloise Keeling: Data Curation Magdalena Martinez: Formal Analysis Jörn Lakowski: Conceptualisation, Methodology, Supervision, Writing- review & editing revision Andrew Lotery: Conceptualisation, Funding acquisition, Supervision, Writing- Review & Editing

