## Supplemental Figure 1 for "TSPAN10 Loss-of-Function in hESC-Derived RPE Reveals Critical Roles in Pigmentation, Polarity, and Antioxidant Defence"

Figure S‑1 Sanger sequence traces of both TSPAN10 mutants in comparison to WT at Exon 2. Related to Figure 1

(A) **Sanger sequence traces for hESC clone A2** (top sequence) vs WT (bottom sequence). 7 bp deletion from 194 bp in clone A2 outlined in red.
(B) **Sanger sequence traces for hESC clone A12** (top sequence) vs WT (bottom sequence). 1 bp insertion of A at 197 bp in clone A12 outlined in red.

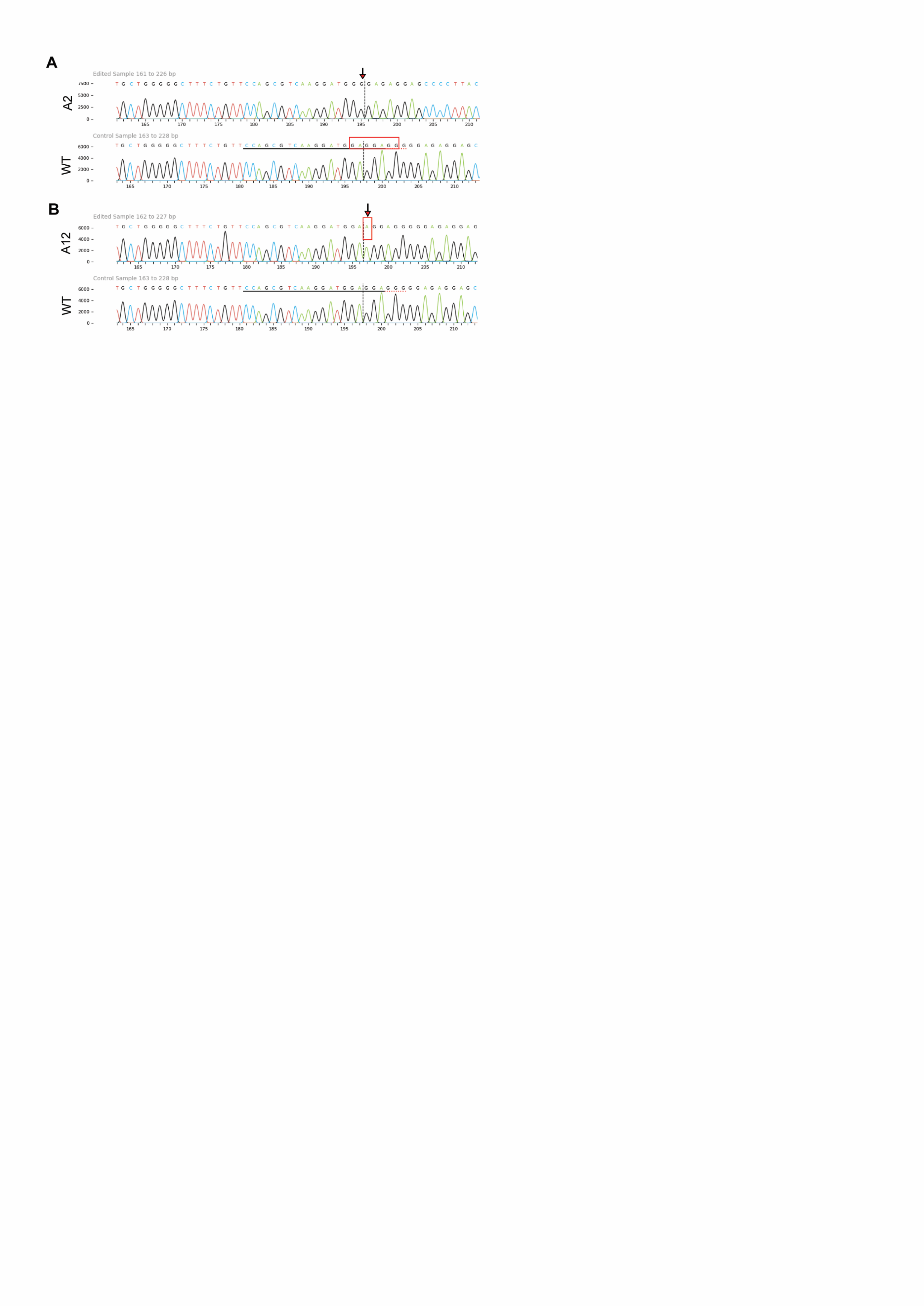
