## Supplemental Figure 2 for "TSPAN10 Loss-of-Function in hESC-Derived RPE Reveals Critical Roles in Pigmentation, Polarity, and Antioxidant Defence"

Figure S‑2 Chromatograms of off - target loci sequencing for TSPAN10 mutants and WT cells

Off-target analysis. Related to Figure 1

The first 9 off-target sites predicted for TSPAN10 sgRNA were amplified from genomic DNA by PCR and sequenced with Sanger sequencing.

Sequences are of the predicted sgRNA off-target locus + PAM site. The PAM is underlined in red.

Off target 4 in TSPAN10 k.o A2 shows a 7bp deletion, not predicted to cause significant or pathogenic changes.

A substitution of C->G is seen at Off Target 3 in both TSPAN10 mutants and WT cells, not predicted to cause significant or pathogenic changes.

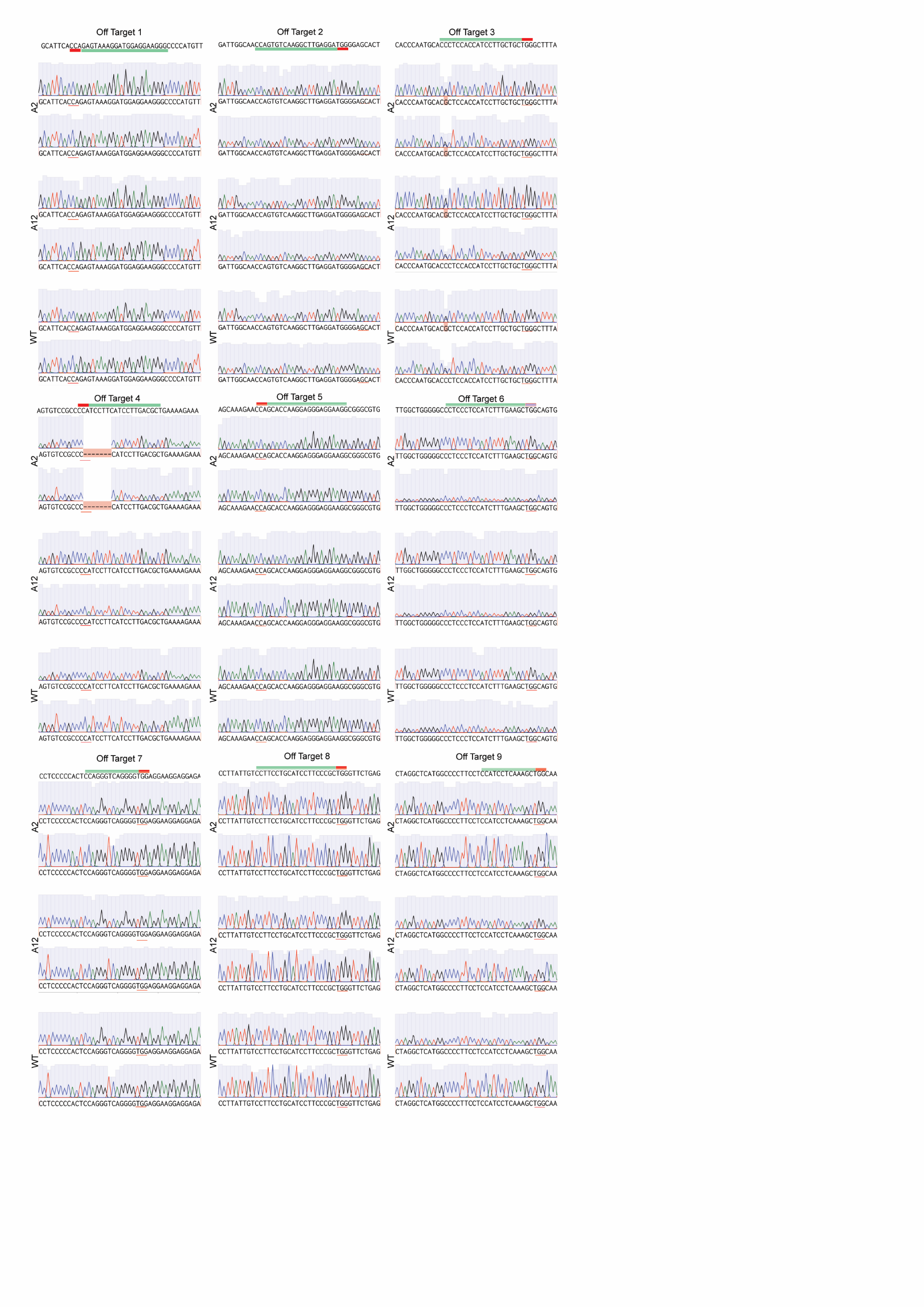
