## Supplemental Figure 4 for "TSPAN10 Loss-of-Function in hESC-Derived RPE Reveals Critical Roles in Pigmentation, Polarity, and Antioxidant Defence"

Figure S‑4 Comparative genome hybridization (array-CGH) report for A12. Related to Figure 1

Array comparative genome hybridization showed no new pathogenic mutations.

No new copy number variants were detected in the A12 hESC line compared to the parental Mastershef7 line (passage 6). Benign CNVs detected in the genetic background of Mastershef7 in our analysis involve chr8 and 15. In addition, we detected one variant of unknown clinical significance (chr7).

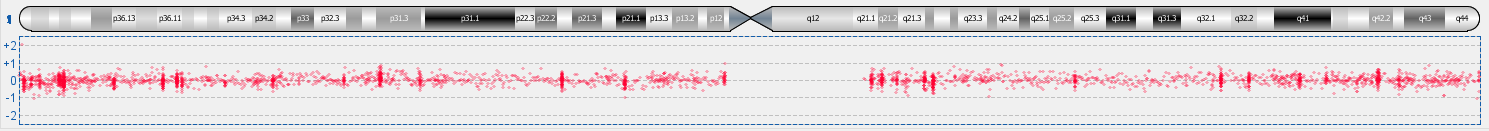

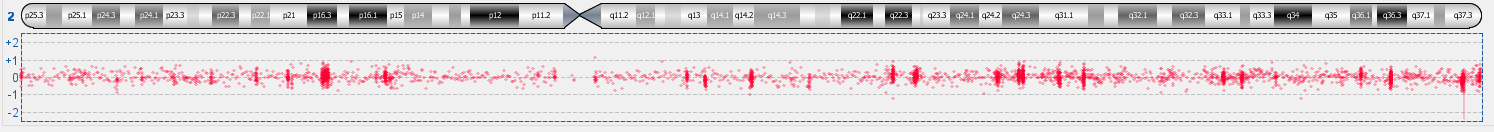

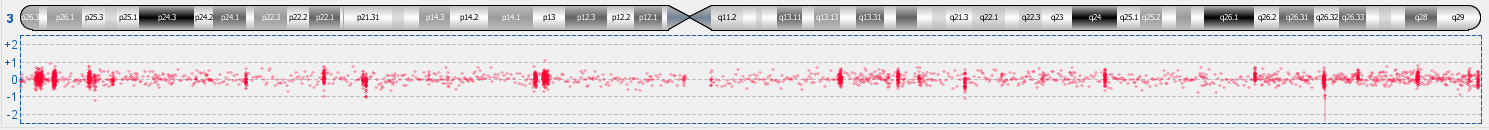

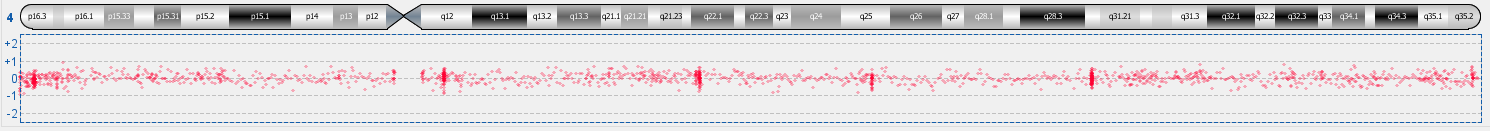

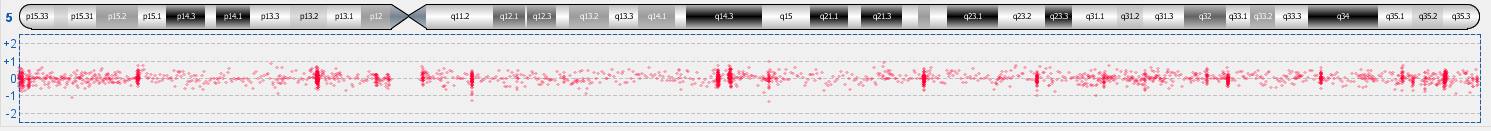

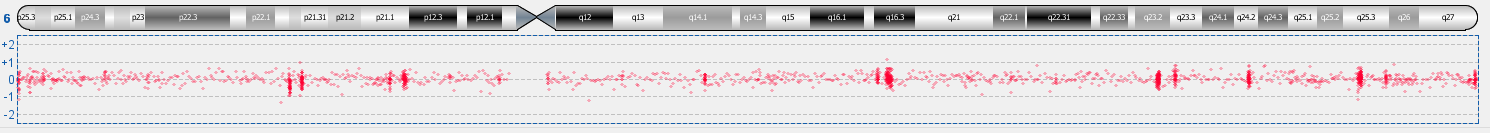

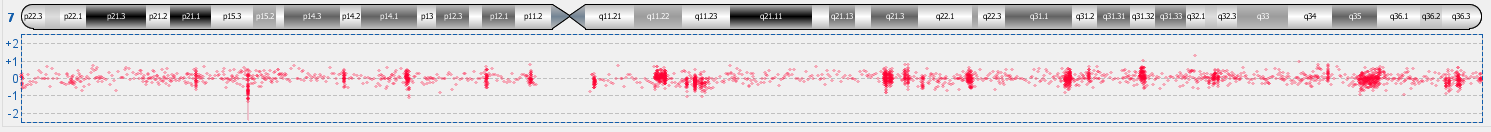

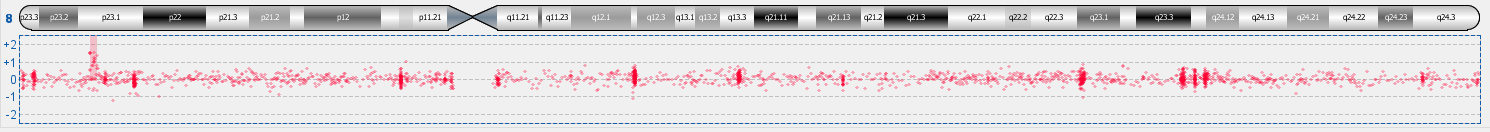

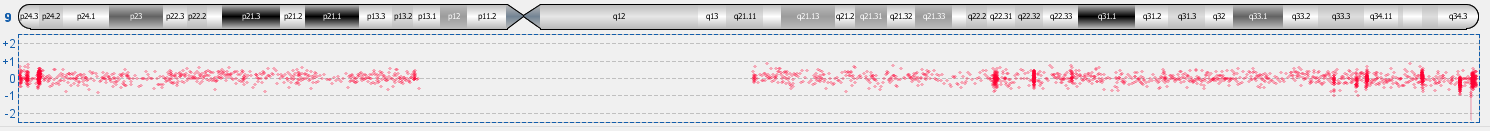

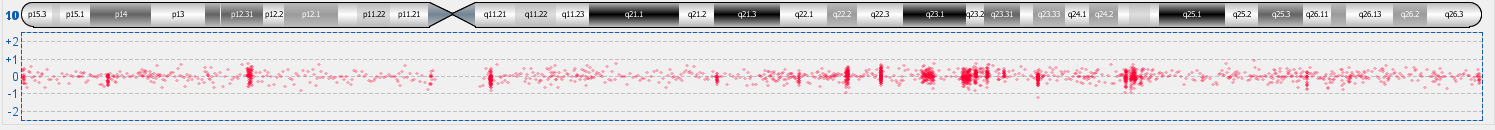

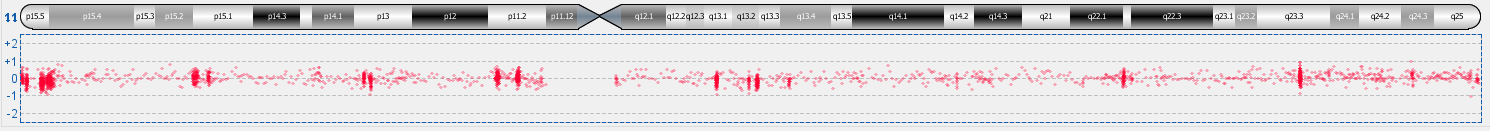

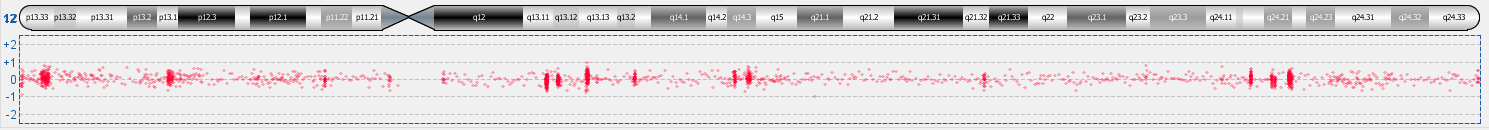

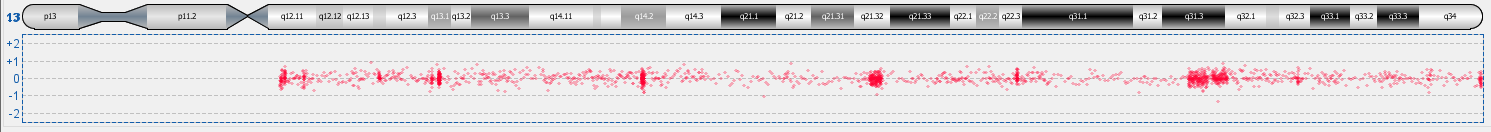

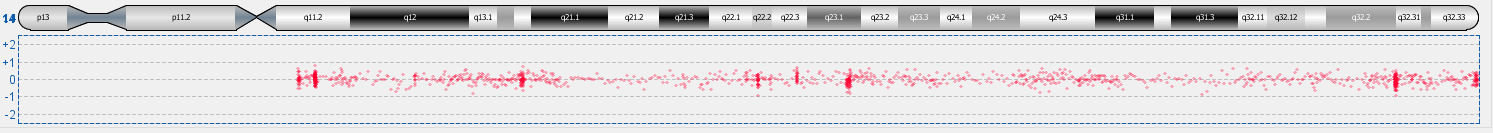

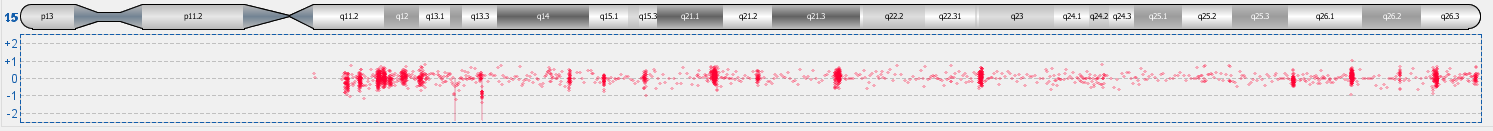

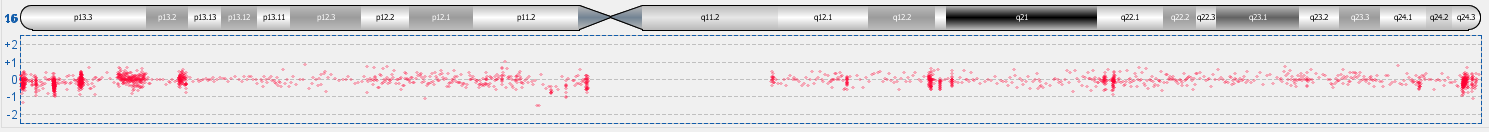

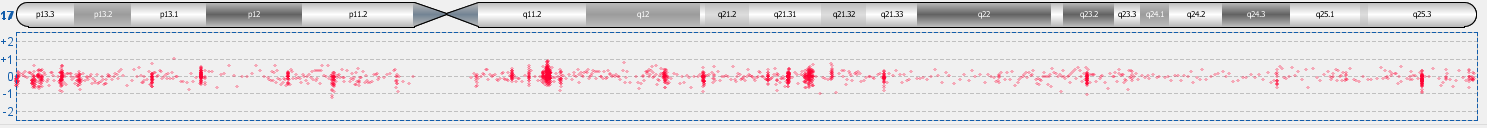

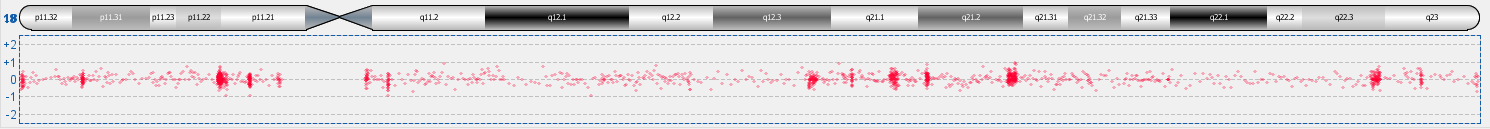

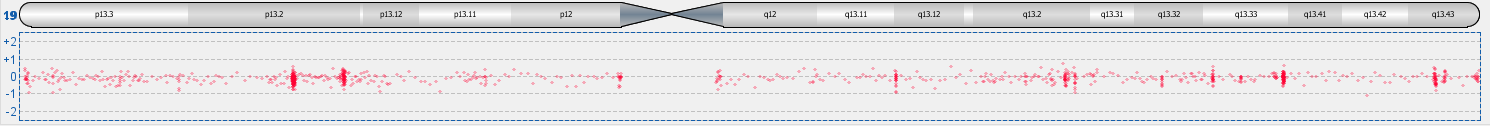

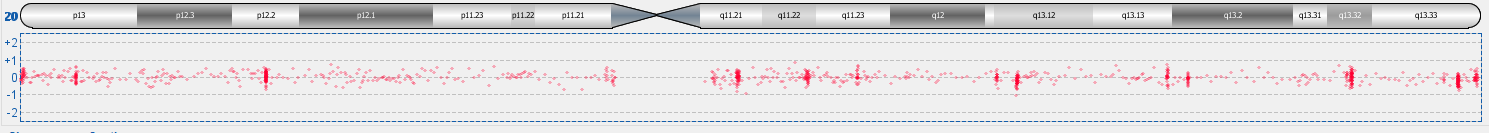

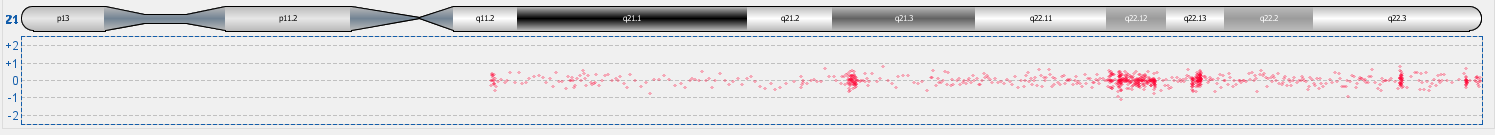

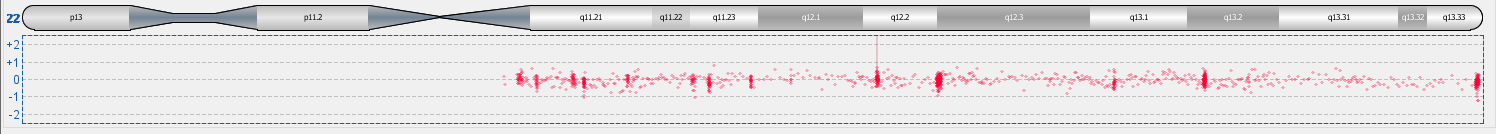

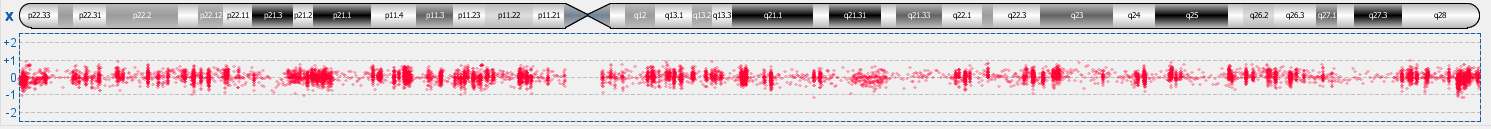

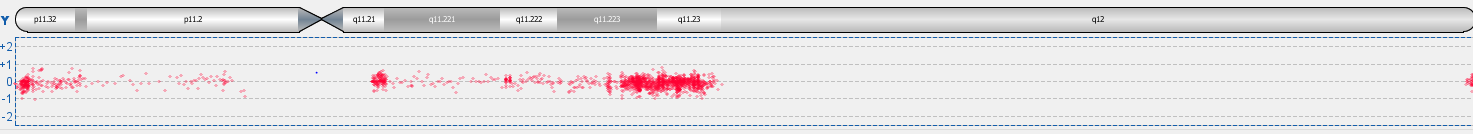
