## Supplemental Figure 5 for "TSPAN10 Loss-of-Function in hESC-Derived RPE Reveals Critical Roles in Pigmentation, Polarity, and Antioxidant Defence"

Figure S‑5 Immunocytochemistry for Na+/K+ ATPase in hESC-RPE

Related to Figure 2. Confocal z-stack images of immunofluorescent labelling for the RPE marker Na^+^/K^+^-ATPase (green) and nuclear stain DAPI (blue) of hESC-RPE cultured on transwells (n=3 for each clone).

Predominantly apical localisation of Na^+^/K^+^-ATPase pump in WT cells.

Apical localisation is lost in A2 and A12.

Scale bars represent 25µm.

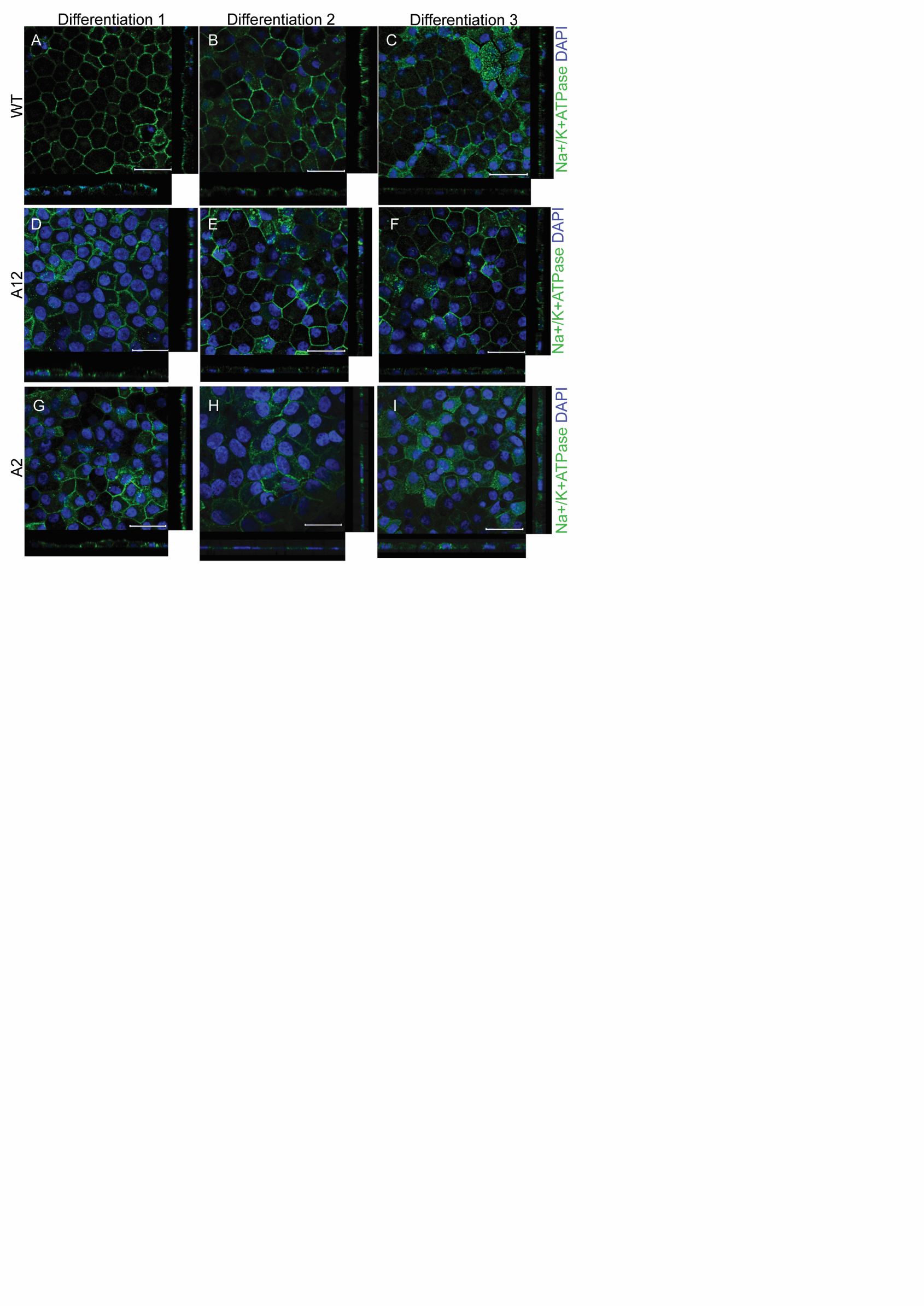
