## Supplemental Figure 6 for "TSPAN10 Loss-of-Function in hESC-Derived RPE Reveals Critical Roles in Pigmentation, Polarity, and Antioxidant Defence"

Figure S‑6 Assessment of TSPAN10 and Rab7 co-localisation in wild-type hESC-RPE.

Ai-Aiv: Confocal micrographs to assess co-localisation of Rab7 (green) with TSPAN10 (red) in WT hESC-RPE. DAPI nuclear stain shown in grey.

B Scatterplot of green (Rab7) and red (TSPAN10) pixel intensities for panel A. The results indicate 95.5% of red (TSPAN10) is colocalised with green (Rab7) and 88.0% of green (Rab7) is colocalised with red (TSPAN10). Intensities below the threshold determined by the Costes et al. algorithm for the x and y axis are not shown (black).

C: Quantification of Rab7 and TSPAN10 co-localisation in WT hESC-RPE. Costes overlap coefficients M1 and M2 indicate the percentage of red (TSPAN10) colocalised with green (Rab7) and green colocalised with red respectively. The results show 89.0 ± 2.0% of Rab7 is colocalised with TSPAN10. Measurements are presented for 3 differentiations. Quantification was performed in the Volocity software suite (PerkinElmer, UK). Performed in triplicate for each differentiation, with three fields of view per differentiation. Scale bars 25 μm

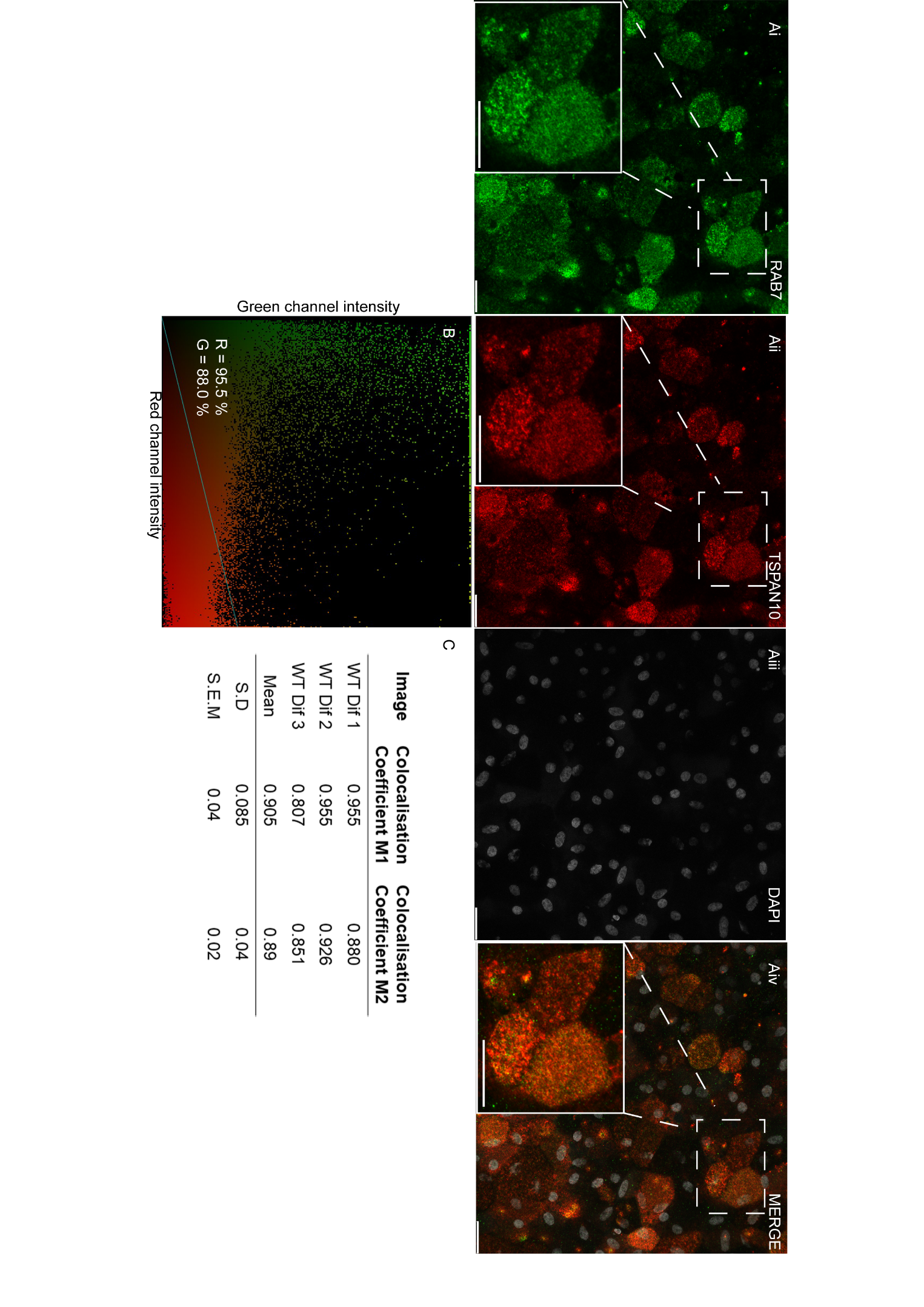
