## Supplemental Figure 7 for "TSPAN10 Loss-of-Function in hESC-Derived RPE Reveals Critical Roles in Pigmentation, Polarity, and Antioxidant Defence"

Figure S‑7 Fluoresence Activated Cell Sorting of hESC-RPE cells for CD140b. Related to Figure 3

(A) hESC-RPEs gated through the FSC-A vs SSC-A plot were further interrogated by the ratios of height to width in forward scatter and side scatter, in addition to their ability to uptake the live/dead cell marker eFluor 506. Live CD140b^+^ hESC-RPE cells were then selected for cell sorting as shown in B.
(B) Comparison of CD140b^+^ cells between WT and TSPAN10 k.o cells shows an increase in CD140b^+^ cells in TSPAN10 k.o cell lines in comparison to WT.
(C) Heat killed cell control. All cells are gated as positive for the live/dead marker.

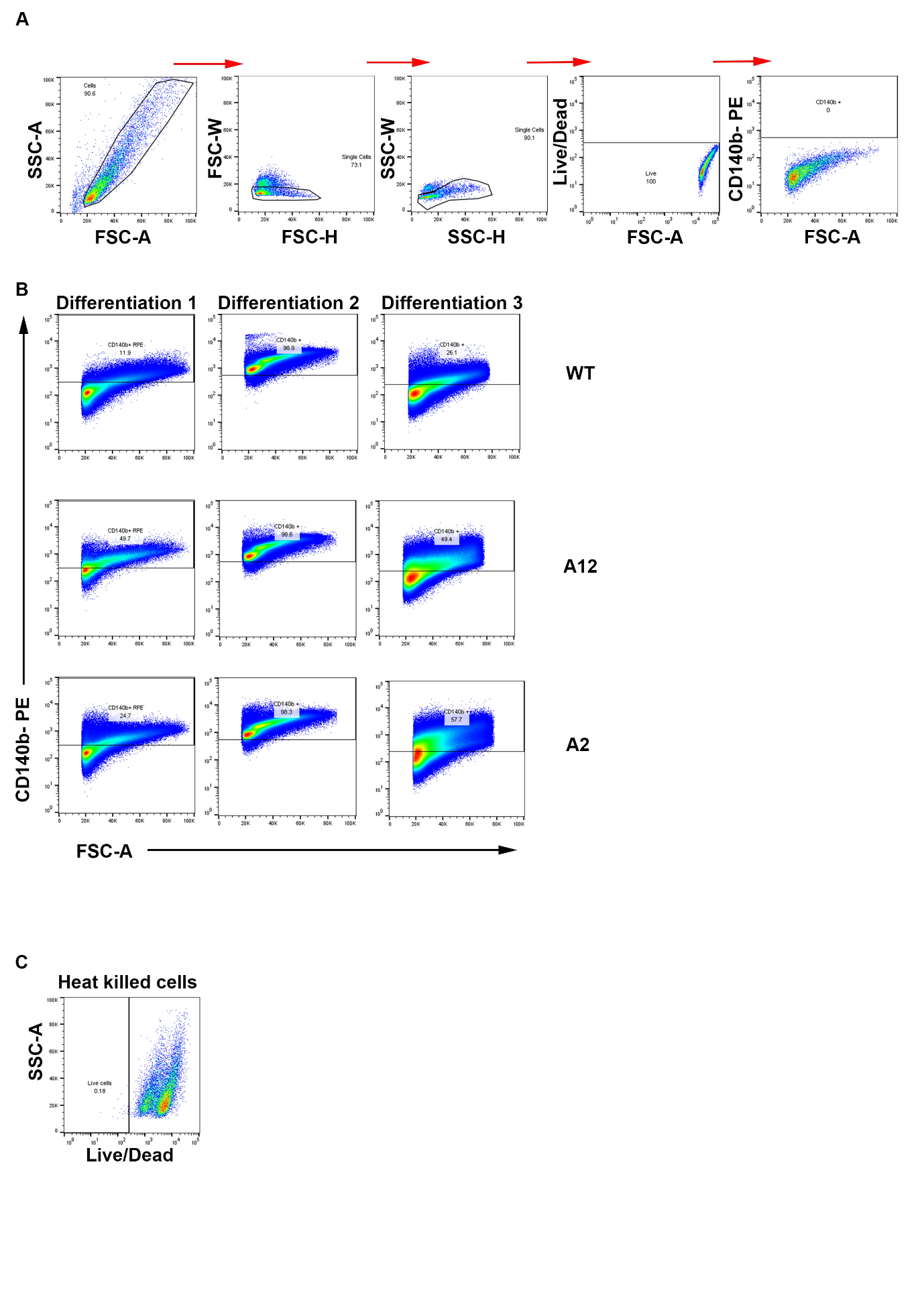
