## Supplemental Figure 8 for "TSPAN10 Loss-of-Function in hESC-Derived RPE Reveals Critical Roles in Pigmentation, Polarity, and Antioxidant Defence"

Figure S‑8 Thresholding of brown pigment in brightfield images of hESC-RPE. Related to Figure 3

Depiction of the steps taken to quantify the level of pigmentation in hESC-RPE cultures.

Initial images show the splitting of channels into red, blue and green and thresholding of the red channel to remove transwell pores to prevent false positives. Subsequent images show the conversion of WT, A12 and A2 brightfield images into binary images where brown pigment has been thresholded (pigment depicted as white, no pigment depicted as black).

Scale bars represent 100 µm.

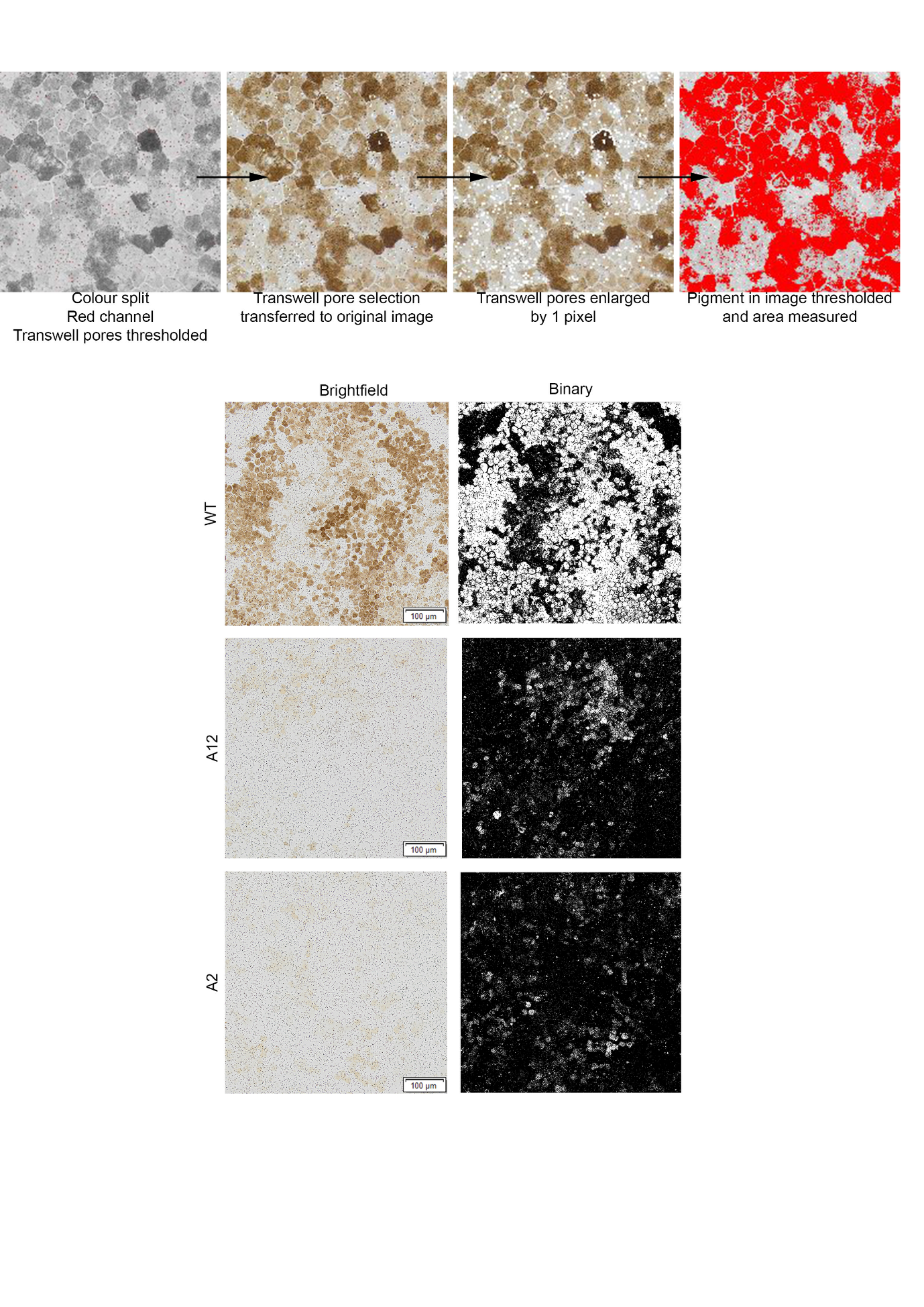
