## Supplemental Figure 9 for "TSPAN10 Loss-of-Function in hESC-Derived RPE Reveals Critical Roles in Pigmentation, Polarity, and Antioxidant Defence"

Figure S-9 Summary of TSPAN10 in the RPE

Schematic depicting the various effects of knocking out TSPAN10 using CRISPR/Cas9 in hESC-RPE. Our results show an effect on pigmentation, tyrosinase expression and activity. Catalase expression and activity, RPE cell polarisation and cell height. Created in BioRender. Kaye, R. (2026) <https://BioRender.com/6mryu4x>

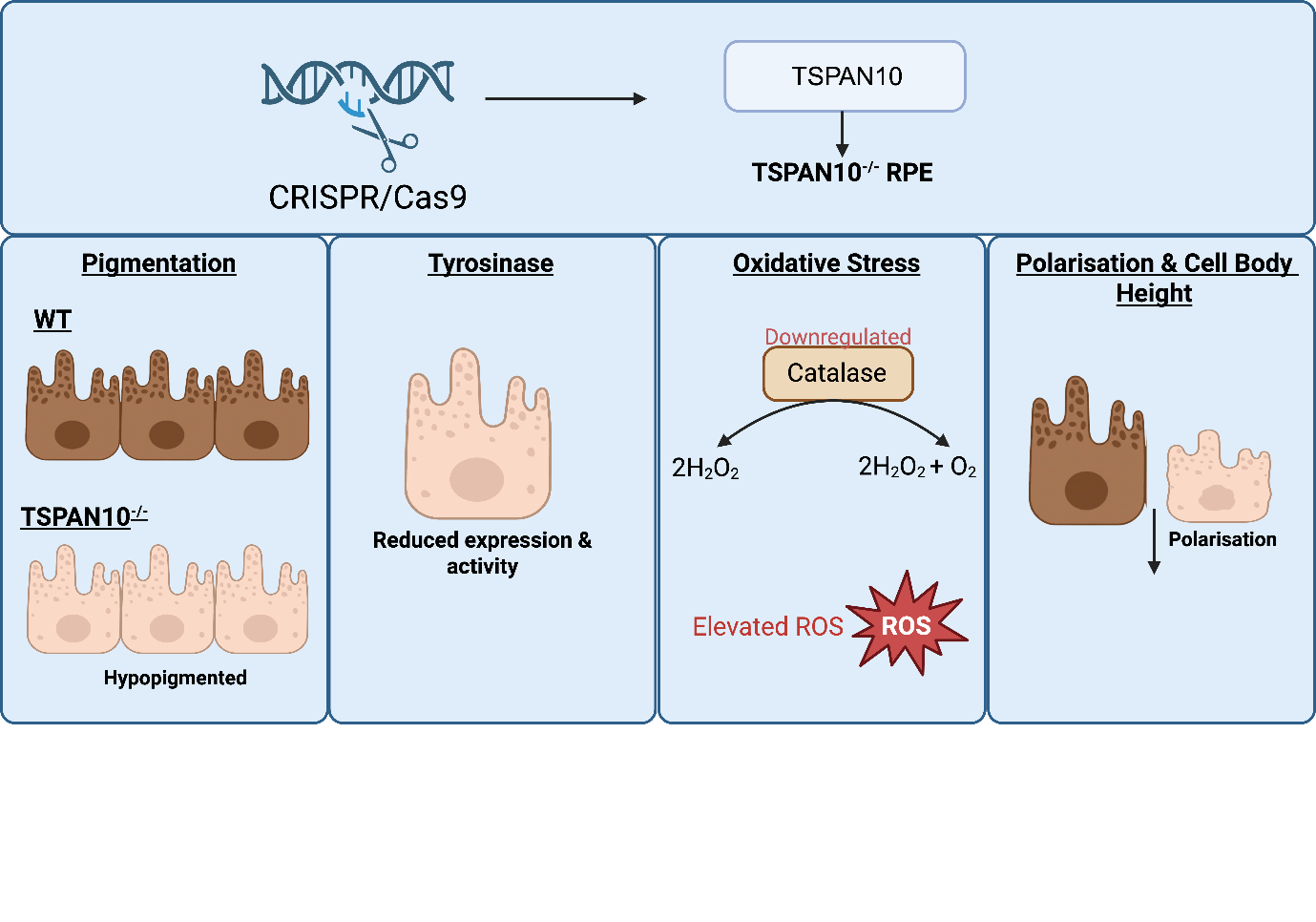
