## Supplemental Figure 10 for "TSPAN10 Loss-of-Function in hESC-Derived RPE Reveals Critical Roles in Pigmentation, Polarity, and Antioxidant Defence"

Figure S-10 Immunocytochemistry of hESCs for pluripotency markers

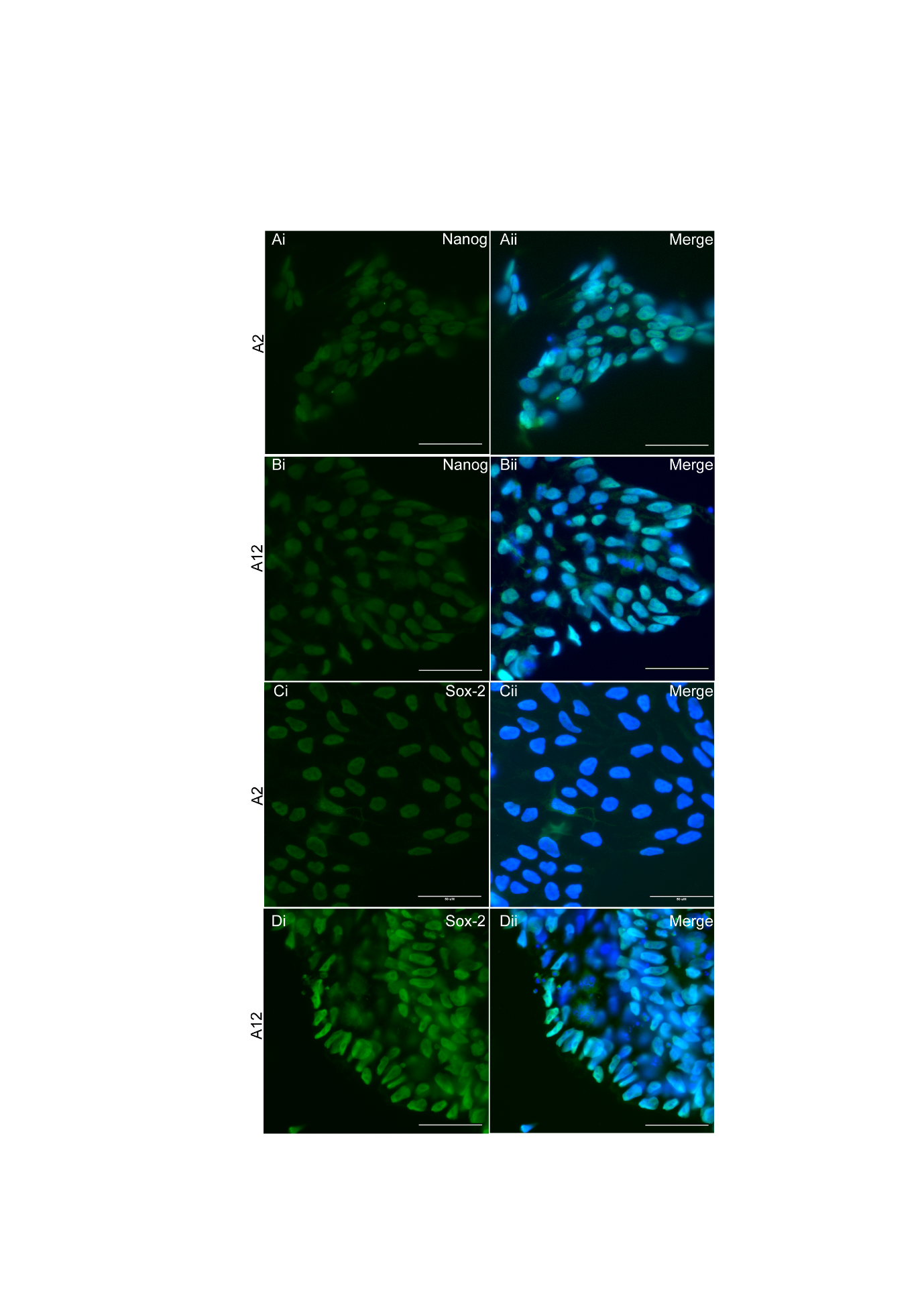

Immunofluorescence images illustrating the expression of pluripotency markers in hESCs. Nuclei are counterstained with DAPI (blue).

Ai-Aii. A2 express the pluripotency marker Nanog

Bi-Bii. A12 express the pluripotency marker Nanog.

Ci-Cii. A2 express the pluripotency marker Sox-2

Di-Dii. A12 express the pluripotency marker Sox-2.

Scale bars represent 50 µm.
