## Supplemental Table 1 for "TSPAN10 Loss-of-Function in hESC-Derived RPE Reveals Critical Roles in Pigmentation, Polarity, and Antioxidant Defence"

| Off-target DNA sequence + PAM | Chromosome | Position | Direction | Mismatches | Off Target |
| --- | --- | --- | --- | --- | --- |
| CCAGCACCAAGGAGGGAGGAAGG | chr8 | 1914086 | + | 3 | 5 |
| TCAGCGTCAAGGATGAAGGATGG | chr5 | 142223136 | - | 2 | 4 |
| CCAGCTTTGAGGATGGAGGAAGG | chr20 | 59505209 | - | 3 | 9 |
| CCAGTGTCAAGGCTTGAGGATGG | chr2 | 148592319 | + | 3 | 2 |
| CCAGAGTAAAGGATGGAGGAAGG | chr2 | 168339209 | + | 2 | 1 |
| CCAGGGTCAGGGGTGG | chr12 | 89392162 | + | 3 | 7 |
| CCAGCAGCAAGGATGGTGGAGGG | chr4 | 189658739 | - | 3 | 3 |
| CCAGCGGGAAGGATGCAGGAAGG | chr17 | 14974956 | - | 3 | 8 |
| CCAGCTTCAAAGATGGAGGGAGG | chr10 | 3180569 | - | 3 | 6 |

**Table S-1 Top nine predicted off target loci.** Related to Figure 1

PAM is underlined; mismatches between the sgRNA on-target and off-target sequences are shown in red; the chromosome and position at which the off-target site is found are reported in addition to the strand. The column off target refers to the name given to each specific locus for alignment with Figure S1.
